# Mapping Along-Tract White Matter Microstructural Differences in Autism

**DOI:** 10.1101/2025.03.21.644498

**Authors:** Gaon S. Kim, Bramsh Q. Chandio, Sebastian M. Benavidez, Yixue Feng, Paul M. Thompson, Katherine E. Lawrence

## Abstract

Previous diffusion magnetic resonance imaging (dMRI) research has indicated altered white matter microstructure in autism, but the implicated regions are highly inconsistent across studies. Such prior work has largely used conventional dMRI analysis methods, including the traditional microstructure model, based on diffusion tensor imaging (DTI). However, these methods are limited in their ability to precisely map microstructural differences and accurately resolve complex fiber configurations. In our study, we investigated white matter microstructure alterations in autism using the refined along-tract analytic approach, BUndle ANalytics (BUAN), and an advanced microstructure model, the tensor distribution function (TDF). We analyzed dMRI data from 365 autistic and neurotypical participants (5-24 years; 34% female) from 10 cohorts to examine commissural and association tracts. Autism was associated with lower fractional anisotropy and higher diffusivity in localized portions of nearly every commissural and association tract examined; these tracts inter-connected a wide range of brain regions, including frontal, temporal, parietal, and occipital. Taken together, BUAN and TDF allow robust and spatially precise mapping of microstructural properties in autism. Our findings rigorously demonstrate that white matter microstructure alterations in autism may be greater within specific regions of individual tracts, and that the implicated tracts are distributed across the brain.

## 1. INTRODUCTION

Autism is a heterogeneous neurodevelopmental condition that affects 1 in 36 children in the United States and an estimated 1 in 100 children worldwide (Zeidan et al. 2022; Maenner 2023). It is clinically defined by a range of social communication challenges and the presence of restricted and repetitive behaviors and interests (Lord et al. 2020; Al-Beltagi 2021; American Psychiatric Association 2022; Hirota and King 2023; Maenner 2023). Studies using structural and functional magnetic resonance imaging (MRI) have consistently linked autism to gross structural and functional alterations in the brain, suggesting the promise of neuroimaging for improving our understanding of variability in this neurodevelopmental condition (Cardinale et al. 2013; Van Rooij et al. 2018; Hong et al. 2019; Lawrence et al. 2019; Postema et al. 2019; Banker et al. 2021; Lawrence et al. 2022; Sha et al. 2022; Duan et al. 2024). Diffusion MRI (dMRI) allows us to specifically examine brain microstructure, including fine-scale properties of the white matter and neural pathways (Le Bihan 2003; Martinez-Heras et al. 2021; Huang et al. 2022). The majority of dMRI research in autism to date has used the traditional reconstruction model, based on diffusion tensor imaging (DTI), together with conventional analytic methods, such as voxel-based analysis (VBA), tract-based spatial statistics (TBSS), or tractography by averaging signals across large white matter tracts (Basser et al. 1994; Travers et al. 2012; Aoki et al. 2013; Ameis and Catani 2015; Di et al. 2018; Fitzgerald et al. 2019; Zhao et al. 2022). These studies have overall shown altered white matter microstructure in autism compared to neurotypical controls, but the white matter regions reported as affected are highly inconsistent across studies (Di et al. 2018; Fitzgerald et al. 2019; Zhao et al. 2022).

A recent dMRI meta-analysis on autism using VBA or TBSS studies examined fractional anisotropy differences based on the DTI model (FA^DTI^) and reported that the left splenium of the corpus callosum and right cerebral peduncle exhibited lower FA^DTI^ in autism, which may reflect reduced white matter integrity (Di et al. 2018). Another VBA/TBSS meta-analysis that focused on neurodevelopmental disorders and two DTI metrics – FA^DTI^ and mean diffusivity (MD) – revealed that autism is associated with lower FA^DTI^ in the genu and splenium of corpus callosum, as well as higher MD in the posterior thalamic radiation (Zhao et al. 2022). Building on these meta-analyses, more recent TBSS studies have similarly reported lower FA^DTI^ in autism compared to neurotypical controls (Haigh et al. 2020; Kilroy et al. 2022; Li et al. 2022). However, the affected regions vary across studies: some studies have found significant commissural tracts alterations in autism, whereas others have reported significant alterations in association tracts (Haigh et al. 2020; Kilroy et al. 2022; Li et al. 2022). Although these findings offer valuable insight into white matter differences in autism, there are limitations of VBA and TBSS. VBA results can be affected by poor registration of anatomy across subjects and difficulty mapping results to specific white matter tracts due to the inherent challenges of resolving overlapping tracts when using voxel-based methods (Bach et al. 2014; Zhou et al. 2022). TBSS addresses the registration concerns of VBA, by mapping individual data to a white matter skeleton, but is limited by potential biases and anatomical inaccuracies during the skeletonization step, as well as similar challenges in tract-specific mapping (Travers et al. 2012; Bach et al. 2014).

Tractography is a complementary analytic method to TBSS and VBA that specifically reconstructs white matter tracts as geometrical models in 3D, often represented as a set of 3D curves or streamlines that make up a fiber bundle (Nimsky et al. 2007; Schmahmann et al. 2007; Jbabdi and Johansen-Berg 2011). Prior tractography studies in autism have identified altered white matter properties most consistently in commissural and association tracts, though the tracts implicated are inconsistent (Aoki et al. 2013; Nordahl et al. 2015; Blanken et al. 2017; Hrdlicka et al. 2019; Zhang et al. 2019; Barbeau et al. 2020; Bassell et al. 2020; Kilroy et al. 2022; Weber et al. 2022; Minnigulova et al. 2023; Oblong et al. 2023; Weerasekera et al. 2024). More recent tractography studies that included over 100 autistic participants have primarily focused on whole-brain organizational properties by using graph theory or incorporating the tractography features into multimodal imaging models (Zhang et al. 2022; Oblong et al. 2023; Weber et al. 2023; Park et al. 2024). These studies overall found atypical structure or structure-function coupling in autism, but the analytic methods used by such work provided limited anatomical specificity of the reported white matter differences (Zhang et al. 2022; Oblong et al. 2023; Weber et al. 2023; Park et al. 2024). However, contextualizing such white matter microstructure alterations by identifying localized patterns of differences will be necessary to comprehensively understand brain circuitry alterations in autism.

To help address the limitations of existing dMRI studies in autism, advanced tractometry techniques such as BUndle ANalytics (BUAN) offer greater spatial precision when characterizing white matter differences. Specifically, BUAN allows for the detailed analysis of microstructure along the entire length of a tract by mapping localized microstructure alterations for each examined tract (Garyfallidis et al. 2014; Chandio et al. 2020). BUAN has successfully been applied to a range of scientific questions, yielding nuanced findings on white matter microstructure in Parkinson’s disease, Alzheimer’s disease, bipolar disorder, and normative aging (Garyfallidis et al. 2014; Chandio et al. 2020; Chandio et al. 2022; Schilling et al. 2022; Nabulsi et al. 2023; Chandio et al. 2025). Such studies applying BUAN to study other brain-based disorders have consistently revealed spatially varying or localized alterations within white matter tracts that may be obscured when using conventional analytic methods that average measures across entire tracts. Notably, one such study found that even the directionality of microstructural alterations can vary within different portions of the same tract, demonstrating the importance of considering fine-grained microstructural differences to obtain a complete understanding of white matter alterations (Nabulsi et al. 2023). Yet, no autism studies to date have completed such spatially-specific microstructure analyses along the length of white matter tracts.

Most autism dMRI studies have used the traditional white matter model, DTI, which has the key limitation that it cannot account for complex white matter configurations: crossing fibers in different three dimensional directions cannot be modeled or visualized using DTI due to the strict Gaussianity of the model, which relies on a single tensor (Basser et al. 1994; Mori and Zhang 2006; Alexander et al. 2007; Jones 2008; Leow et al. 2009; Soares et al. 2013; Nir et al. 2017). This limitation is addressed by advanced dMRI models, including the advanced single-shell model, the tensor distribution function (TDF). TDF overcomes this limitation by fitting a continuous distribution of tensors to the diffusion signal at each location in the brain, allowing for a more accurate representation of complex fiber orientations within each voxel (Leow et al. 2009; Nir et al. 2017). It uses the least-squares principle and gradient descent algorithms to determine the most suitable TDF for the voxel. TDF has been shown to be more sensitive than DTI in previous studies of Alzheimer’s disease, cognitive impairment, and normative development and aging (Nir et al. 2017; Zavaliangos-Petropulu et al. 2019; Lawrence et al. 2021; Benavidez et al. 2024). However, the TDF model has not yet been used to investigate white matter microstructure in autism.

Here, we rigorously examined white matter microstructure differences in autism by applying the refined tractometry method BUAN and the advanced microstructure model TDF along with traditional DTI microstructural metrics, to a large sample of 365 participants from 10 cohorts (Leow et al. 2009; Aoki et al. 2013; Nir et al. 2017; Hrdlicka et al. 2019; Barbeau et al. 2020; Chandio et al. 2020; Kilroy et al. 2022; Olivé et al. 2022). We hypothesized that FA would be lower in the autism group across commissural and association tracts and that the TDF microstructure model would capture white matter differences in autism more sensitively than DTI (Aoki et al. 2013; Nir et al. 2017; Hrdlicka et al. 2019; Kilroy et al. 2022). Based on previous BUAN studies in other brain-based disorders, we also expected microstructural alterations to vary along the length of the tract (Chandio et al. 2020; Chandio et al. 2022; Schilling et al. 2022; Nabulsi et al. 2023). To the best of our knowledge, this large autism dMRI study is also the first study to investigate along-tract microstructural alterations in autism.

## 2. METHODS

### 2.1. Participants and MRI Acquisition

Our final sample consisted of 365 participants between the ages of 5 to 24 years (age: 13.6 ± 3.6 years; 34.0% female). This included 195 autistic individuals (age: 13.4 ± 3.7 years; 34.4% female) and 170 neurotypical controls (age: 13.9 ± 3.3 years; 33.5% female) (**Table 1**). Complete demographic characteristics are summarized in **Table 1**. Data in this sample were drawn from 10 cohorts (sites/scanners), including seven from the NIMH Data Archive (NDA) and three from the Autism Brain Imaging Data Exchange (ABIDE) (**Supplementary Table 1**) (Di Martino et al. 2017). More details on participant recruitment, study inclusion criteria, and scanning protocols are described elsewhere (Lee et al. 2007; Di Martino et al. 2017; Irimia et al. 2017; Jack et al. 2021). Briefly, all dMRI scans were single-shell acquisitions collected on 3T scanners; cohort-specific scanner descriptions and acquisition protocols are provided in **Supplementary Table 2**. To be included in the current analyses, participants from each cohort were required to have a complete dMRI scan and complete information for diagnosis, age, sex, and full-scale intelligence quotient (FSIQ). Subjects were excluded if they had poor quality dMRI data, including excessive head motion (see *2.2 dMRI Data Analysis*). As we were specifically interested in the developmental period from childhood through emerging adulthood, participants were also excluded from the current analyses if they were over 24 years old. For cohorts that included any siblings or any longitudinal scans, we randomly selected one sibling or timepoint per family or participant to ensure independence of data samples in the overall statistical model. When contrasting the autism and neurotypical groups in our final sample, there were no significant group differences in age (*p*=0.11) or sex (*p*=0.86). As expected, FSIQ was higher in neurotypical participants, on average (*p*<0.001), and mean relative head motion was higher in autistic participants, on average (*p*=0.008). We thus included FSIQ and mean relative head motion as covariates in our primary and/or supplemental analyses (see *2.3 Statistical Analyses*).

**Table 1.** Participant demographics.

|  | Autism<br>(N = 195) | Neurotypical<br>(N = 170) | Overall<br>(N = 365) | P-value |
| --- | --- | --- | --- | --- |
| **Sex** |  |  |  |  |
| Male/Female | 128 / 67 | 113 / 57 | 241 / 124 | $p = 0.86$ |
| **Age (years)** |  |  |  |  |
| Mean (SD) | 13.4 ($\pm 3.74$) | 13.9 ($\pm 3.29$) | 13.6 ($\pm 3.55$) | $p = 0.11$ |
| **FSIQ** |  |  |  |  |
| Mean (SD) | 101.7 ($\pm 18.69$) | 111.6 ($\pm 14.85$) | 106.3 ($\pm 17.69$) | $p < 0.001$ |
| **Head Motion (mm)** |  |  |  |  |
| Mean (SD) | 0.312 ($\pm 0.134$) | 0.274 ($\pm 0.134$) | 0.294 ($\pm 0.135$) | $p = 0.008$ |

FSIQ, full-scale intelligence quotient; SD, standard deviation

### 2.2. dMRI Data Analysis

The dMRI scans were preprocessed using the ENIGMA DTI protocol (http://enigma.ini.usc.edu/protocols/dti-protocols) (Jahanshad et al. 2013). Preprocessing included eddy current and head motion correction (using FSL’s *eddy_openmp*), bias field correction (using MRtrix3’s *dwibiascorrect*), and susceptibility artifact correction (using Synb0-DisCo or FSL’s *flirt*), as well as denoising (using theMarcenko-Pastur principal component analysis algorithm as implemented in DIPY’s *genpca*) when appropriate (Tustison et al. 2010; Jenkinson et al. 2012; Garyfallidis et al. 2014; Schilling et al. 2019; Tournier et al. 2019). After preprocessing, we fit the diffusion models, based on the DTI and TDF formulae. DTI, the conventional modeling approach for dMRI data, fits a single-tensor to dMRI data and typically represents hindered diffusion (Basser et al. 1994; Jones 2008). However, DTI has well-established limitations in capturing complex fiber configurations (Jones 2008; Leow et al. 2009). These limitations are addressed by the TDF model developed by our group, which is a more advanced single-shell model (Jones 2008; Leow et al. 2009; Nir et al. 2017). TDF uses a continuous mixture of tensors to account for multiple underlying fiber populations (Leow et al. 2009; Nir et al. 2017). For each voxel we calculated four DTI metrics – fractional anisotropy (FA^DTI^), mean diffusivity (MD), axial diffusivity (AD), radial diffusivity (RD) – and one TDF metric, an advanced measure of fractional anisotropy (FA^TDF^) (Basser et al. 1994; Jones 2008; Leow et al. 2009; Nir et al. 2017). Among DTI metrics, FA^DTI^ measures the degree of anisotropy and MD represents the overall magnitude of water diffusion (Travers et al. 2012; Soares et al. 2013). AD measures diffusion along the primary axis of the tract and RD measures diffusion perpendicular to it (Soares et al. 2013). The TDF metric, FA^TDF^, measures the degree of directional diffusion, as does FA^DTI^, but accounts for crossing fibers (Leow et al. 2009; Nir et al. 2017).

We analyzed white matter microstructure in autism at a finer anatomical scale using the robust, flexible, and advanced tractography-based approach, BUAN (**Figure 1**) (Chandio et al. 2020). BUAN is publicly available through the Diffusion Imaging in Python (DIPY) library (Garyfallidis et al. 2014; Chandio et al. 2020). Using the BUAN tractometry pipeline, we were able to locate diagnosis differences at specific segments along the length of the individual white matter tract. Unlike traditional tractometry methods, BUAN incorporates all streamlines within a bundle rather than reducing it to a single mean streamline. This differentiates BUAN as it does not simplify a large white matter bundle to a single mean streamline like conventional methods but ensures that the entire bundle’s microstructural profile is captured (Yeatman et al. 2012; Chandio et al. 2020; Nabulsi et al. 2023).

**Figure 1.**
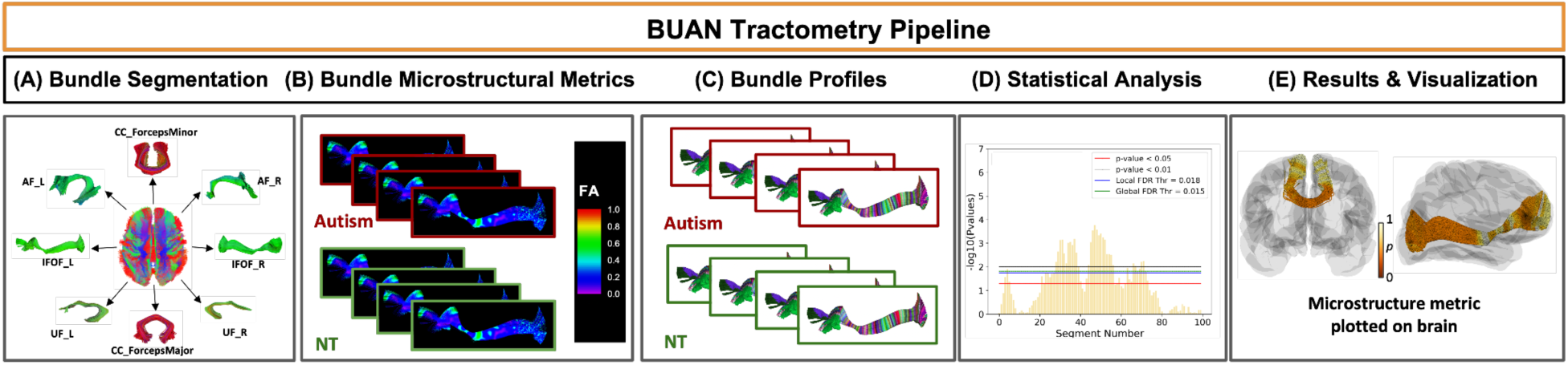
Overview of the BUAN tractometry pipeline. (A) Individual white matter tracts are extracted from a reconstructed whole-brain tractogram. (B) Bundle profiles are created by mapping microstructural measures onto the extracted bundles. (C) Each white matter tract is divided into 100 segments for localized tract investigation. (D) Bundle profiles are analyzed using linear mixed models. *P*-values for the effect of diagnosis are then plotted from these statistical analyses for each microstructural metric at each segment; multiple *p*-value thresholds for significant differences are shown for completeness. (E) These *p*-values are mapped onto the tract in a 3D “glass brain” for visualization.

As in our prior work developing and applying BUAN, whole brain tractograms were generated using the constrained spherical deconvolution (CSD) reconstruction model and local deterministic tractography with the EuDx tracking algorithm (Garyfallidis 2012; Jeurissen et al. 2014; Chandio et al. 2020; Nabulsi et al. 2023). The following parameters were used for our tracking algorithm: fiber tracking started from voxels with FA values over 0.3, seed count of 10, continuing with a step size of 0.5, an angle curvature threshold of 25°, and tracking stopped at a minimum FA value of 0.2. The individual whole-brain tractograms were affinely registered to HCP-842 template tractogram in the MNI space (Garyfallidis et al. 2015; Yeh et al. 2018; Chandio et al. 2020) using streamline-based linear registration (SLR) method. Next, a step known as *RecoBundles* was used to extract 9 individual white matter tracts for each subject using the whole brain tractogram and model bundles from the HCP-842 template included in BUAN (Garyfallidis et al. 2014; Garyfallidis et al. 2018; Yeh et al. 2018; Chandio et al. 2020). A tract profile for each bundle was then obtained, whereby the DTI and TDF metrics were mapped onto all the points of the streamlines in a bundle for each subject (Garyfallidis et al. 2012; Garyfallidis et al. 2014; Chandio et al. 2020). Each tract is divided into 100 segments along its length, with each point on a streamline assigned a segment number based on its closest Euclidean distance to a model bundle’s centroid. Lastly, each segment is analyzed using linear mixed models (LMM), allowing for localized statistical assessments (see *2.3 Statistical Analyses* below); to account for multiple fiber observations within a single subject’s tract, subjects are modeled as a random effect in the LMM framework.

To help ensure data quality, we completed rigorous quality control throughout the processing pipeline, as in our prior work (Benavidez et al. 2024). First, all raw dMRI scans were visually inspected for quality assurance to exclude scans with artifacts. After each step in the dMRI processing pipeline, we also visually inspected 2D images of the processing output. Subjects flagged during the visual inspection underwent detailed visual inspection of their whole brain tractograms for a tractography-specific quality control. Participants with poor-quality data such as tract missingness were excluded prior to conducting group-level analyses.

Here, we focused on 9 commissural and association tracts for which mapping accuracy was validated in the foundational BUAN report (Chandio et al. 2020). This included 3 commissural tracts: the *forceps minor*, *forceps major*, and midbody of the corpus callosum. The 6 association tracts consisted of the arcuate fasciculus (AF), extreme capsule (EMC), inferior fronto-occipital fasciculus (IFOF), inferior longitudinal fasciculus (ILF), middle longitudinal fasciculus (MdLF), and uncinate fasciculus (UF). Bilateral association tracts were extracted separately for the left and right hemispheres, resulting in a total of 15 tracts.

### 2.3. Statistical analyses

Linear mixed models (LMMs) were used to examine differences between the autism and neurotypical groups for each tract segment and metric. Specifically, each white matter microstructural metric, such as FA^DTI^, was set as the dependent variable in the LMM. Diagnosis, age, demeaned age squared, sex, and FSIQ were included as fixed effects, and our analyses focused on the effect of diagnosis; the other variables were included as nuisance covariates.

Additionally, we conducted supplementary analyses that covaried for head motion. As in our prior work, subject and cohort were modeled as nested random effects in all analyses (Chandio et al. 2020; Nabulsi et al. 2023); the former was included to account for the non-independence of different streamlines within a single tract segment for a given subject, and the latter was included to model potential effects of cohort (Chandio et al. 2020; Nabulsi et al. 2023; Chandio et al. 2024).

Correction for multiple comparisons was conducted by controlling the False Discovery Rate (FDR) (Benjamini and Hochberg 1995). Similar to prior work, we visualize results at several *p*-value thresholds for completeness and to allow comparability with other studies: uncorrected *p* < 0.05, uncorrected *p* < 0.01, global FDR-corrected threshold, and local FDR-corrected threshold. Global FDR correction was applied to p-values from all 100 segments pooled across 15 tracts (100 × 15), resulting in a global FDR threshold, while tract-specific FDR correction was applied to p-values across the 100 segments within each individual tract, providing a local FDR threshold (Chandio et al. 2020; Nabulsi et al. 2023). We focus on results obtained at the local FDR threshold as in prior work (Nabulsi et al. 2023); we also confirmed the robustness of our results by using the global FDR threshold. Quantile-Quantile (QQ) plots were then created to assess the statistical impact of including additional cohorts and, separately, to investigate which microstructure metric exhibited the smallest *p*-values for each tract. We created the QQ plots by pooling and visualizing the *p-*values across increasing numbers of cohorts and, separately, per metric for each significant tract (**Figure 2-9, Supplementary Figure 1**) (Chandio et al. 2024). QQ plots offer a way to understand the overall pattern of significance in a *p*-value map, in this case for a tract, as higher departures from the diagonal line can be caused by effects at different significance levels. As a heuristic, we often regard a map as having stronger effects in aggregate if the QQ curve is higher.

**Figure 2.**
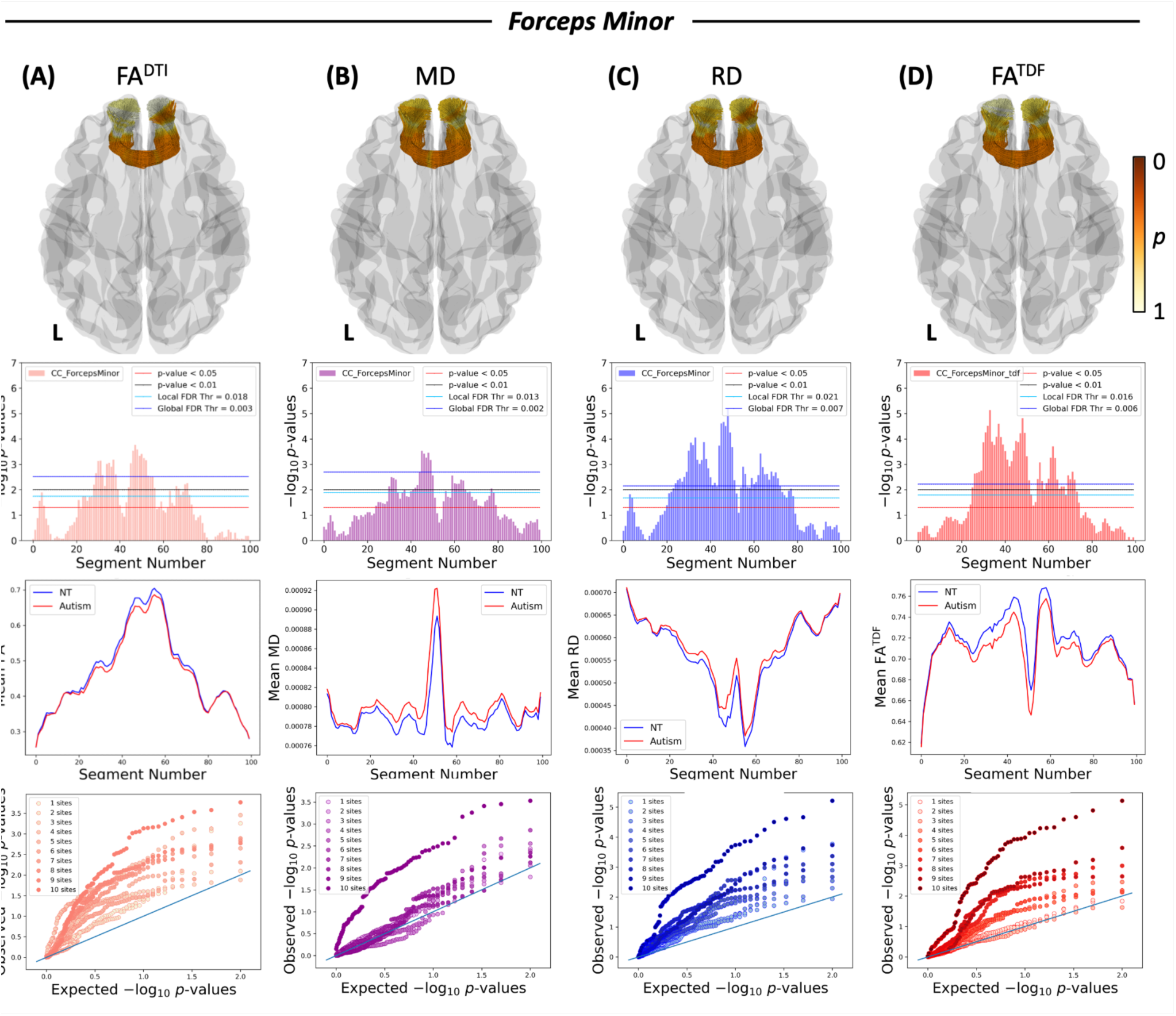
White matter microstructural alterations in the *forceps minor* in autism compared to neurotypical controls. Each column represents a different white matter metric: (A) FA^DTI^, (B) MD, (C) RD, and (D) FA^TDF^. The first row is a 3D representation of the *forceps minor* illustrating the anatomical location and corresponding *p*-values of the corresponding microstructural metric; L indicates the left brain hemisphere and the colorbar shows the *p*-values, where darker orange is a lower *p*-value nearing 0 and lighter yellow is a *p*-value of 1. The second row depicts the negative logarithms of *p*-values for each segment along the tract when contrasting the autism and neurotypical groups; the right hemisphere’s anterior portion of the tract is segment 1, and the segment number increases along the length of the tract up to segment 100 on the left hemisphere’s anterior portion of the tract. The third row depicts mean plots showing the mean microstructural metric for each group within each segment with autism in red and neurotypicals in blue. The fourth row consists of QQ plots that display the effect of adding more cohorts, where including additional cohorts increases sensitivity to white matter differences. FA^DTI^, fractional anisotropy as calculated by the diffusion tensor imaging model; MD, mean diffusivity; RD, radial diffusivity; FA^TDF^, fractional anisotropy as calculated by the tensor distribution function model.

## 3. RESULTS

Microstructural alterations in autism were observed in localized regions of all three commissural tracts examined (**Figure 2-4**). The *forceps minor, forceps major*, and midbody of the corpus callosum all had lower FA^DTI^, as well as higher MD and RD, in the autism group compared to the neurotypical group. No significant differences in AD were detected between groups. Compared to neurotypical controls, FA^TDF^ was lower for all three tracts in autism. Across metrics, the significant microstructure differences observed in the *forceps minor* and midbody of the corpus callosum were specifically localized to the central portion of the tract (**Figure 2-3**). On the other hand, significant differences in the *forceps major* were found in many localized segments across the tract, particularly in the lateral portions (**Figure 3**). These results remained significant with the global FDR threshold. Including additional cohorts provided greater sensitivity when capturing white matter alterations for every commissural tract examined (**Figure 2-4**). When considering the FA metrics derived from our two distinct microstructure models, FA^TDF^ exhibited greater statistical significance overall than FA^DTI^ for all three commissural tracts (**Supplementary Figure 1A-C**).

**Figure 3.**
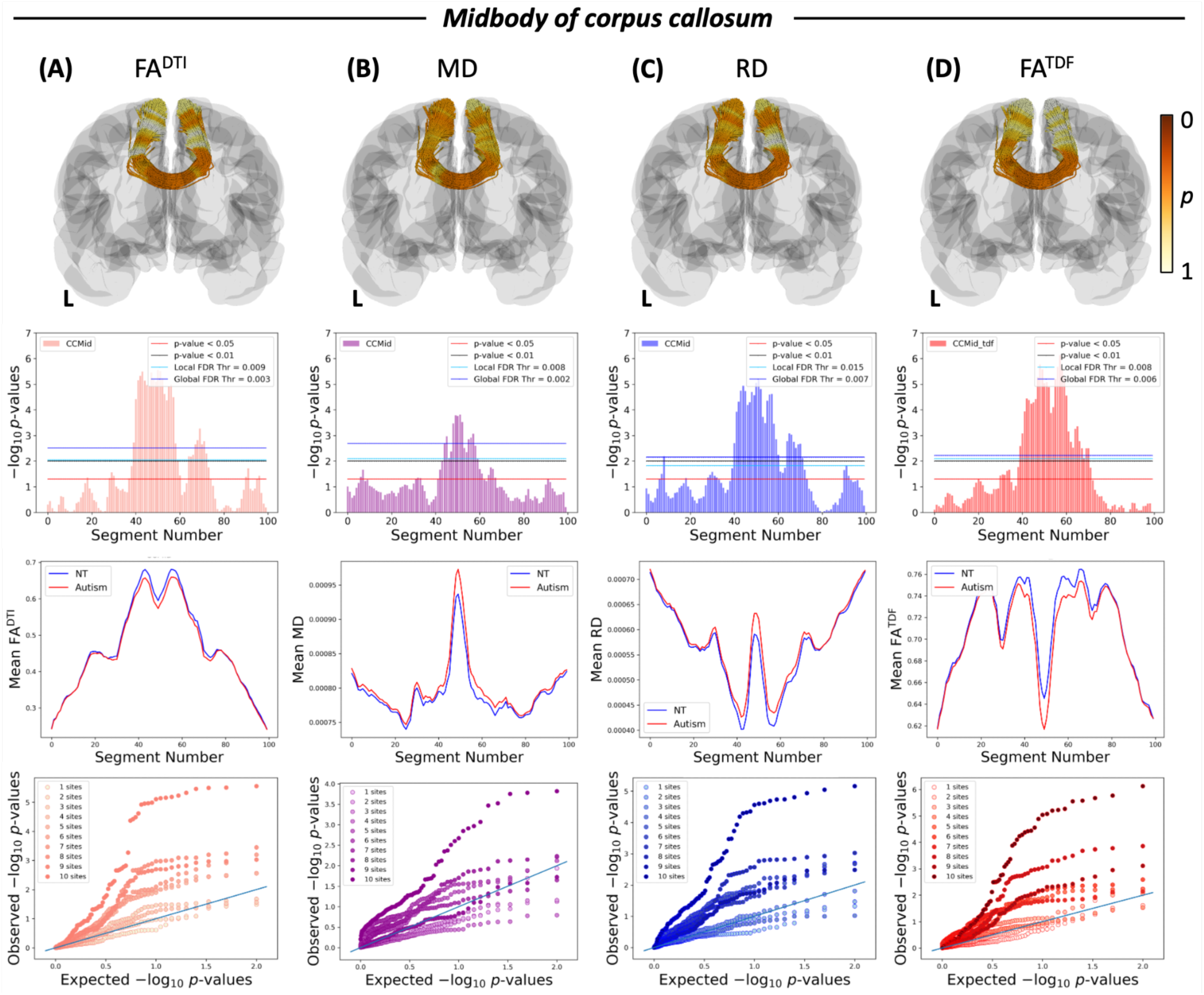
White matter microstructural alterations in the midbody of the corpus callosum in autism compared to neurotypical controls. Each column represents a different white matter metric: (A) FA^DTI^, (B) MD, (C) RD, and (D) FA^TDF^. The first row is a 3D representation of the corpus callosum midbody illustrating the anatomical location and corresponding *p*-values of the corresponding microstructural metric; L indicates the left brain hemisphere and the colorbar shows the *p*-values, where darker orange is a lower *p*-value nearing 0 and lighter yellow is a *p*-value of 1. The second row depicts the negative logarithms of *p*-values for each segment along the tract when contrasting the autism and neurotypical groups; the left hemisphere’s superior portion of the tract is segment 1, and the segment number increases along the length of the tract up to segment 100 on the right hemisphere’s superior portion of the tract. The third row depicts mean plots showing the mean microstructural metric for each group within each segment with autism in red and neurotypicals in blue. The fourth row consists of QQ plots that display the effect of adding more cohorts, where including additional cohorts increases sensitivity to white matter differences. FA^DTI^, fractional anisotropy; MD, mean diffusivity; RD, radial diffusivity; FA^TDF^, tensor distribution function fractional anisotropy.

**Figure 4.**
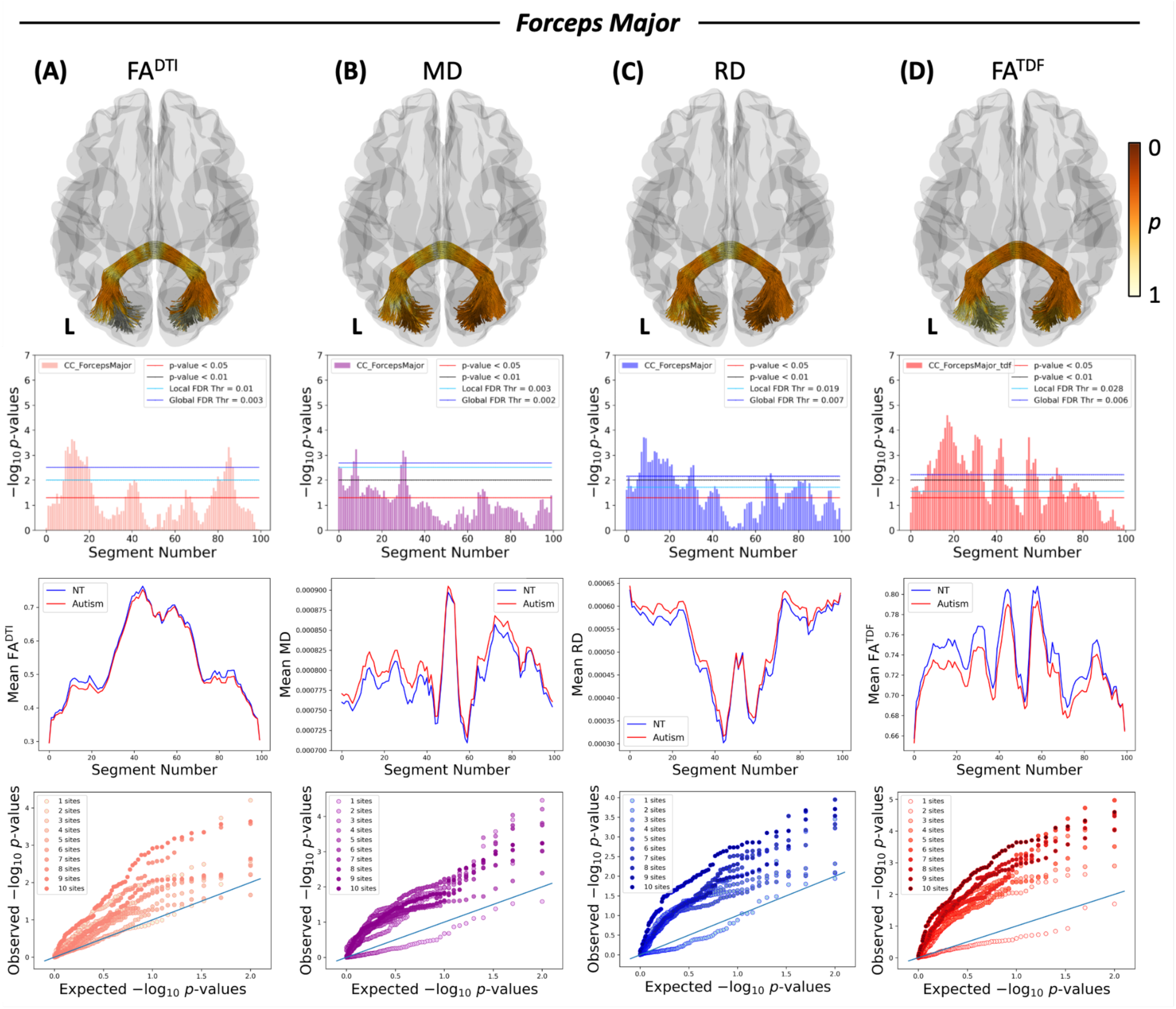
White matter microstructural alterations in the *forceps major* in autism compared to neurotypical controls. Each column represents a different white matter metric: (A) FA^DTI^, (B) MD, (C) RD, and (D) FA^TDF^. The first row is a 3D representation of the *forceps major* illustrating the anatomical location and corresponding *p*-values of the corresponding microstructural metric; L indicates the left brain hemisphere and the colorbar shows the *p*-values, where darker orange is a lower *p*-value nearing 0 and lighter yellow is a *p*-value of 1. The second row depicts the negative logarithms of *p*-values for each segment along the tract when contrasting the autism and neurotypical groups; the right hemisphere’s posterior portion of the tract is segment 1, and the segment number increases along the length of the tract up to segment 100 on the left hemisphere’s posterior portion of the tract. The third row depicts mean plots showing the mean microstructural metric for each group within each segment with autism in red and neurotypicals in blue. The fourth row consists of QQ plots that display the effect of adding more cohorts, where including additional cohorts increases sensitivity to white matter differences. FA^DTI^, fractional anisotropy; MD, mean diffusivity; RD, radial diffusivity; FA^TDF^, tensor distribution function fractional anisotropy.

All six association tracts examined here displayed significant microstructural differences in autism within localized segments of the tract, with the exception of the EMC which did not significantly differ between diagnostic groups (**Figure 4-8**). The AF displayed higher RD in autism than neurotypical controls in both the left and right hemispheres. The right AF also had lower FA^DTI^ and FA^TDF^ in the autism group. Localized differences in the AF were observed in frontal projections of the tract. The right UF showed lower FA^DTI^ and higher RD in autism and, similar to the AF, had localized differences in frontal tract regions. The right IFOF had higher MD and RD in autism compared to neurotypical controls, as well as lower FA^TDF^. Significant differences in the IFOF were localized across metrics in the posterior half of the tract. For the ILF, higher RD and lower FA^TDF^ were found in both the left and right hemispheres in the autism group compared to the neurotypical group. The right ILF additionally had higher MD in the autism group. Microstructure alterations in both the left and right ILF were specific to the posterior and middle portions of the tract. Lastly, the right MdLF had lower FA^DTI^ and FA^TDF^, as well as higher MD, in autism relative to neurotypicals; these differences were localized to the middle portion of the tract. All significant association tracts and metrics were confirmed with the global FDR threshold. For all association tracts examined here that exhibited significant microstructure alterations, including more cohorts increased sensitivity to such differences (**Figure 5-9**). When considering metrics derived from our two distinct microstructure models, DTI and TDF, FA^TDF^ exhibited greater statistical significance in the right IFOF (**Supplementary Figure 1E**). For the right UF and right AF, the most sensitive metric was FA^DTI^. For the right MdLF and right ILF, MD displayed the greatest significance (**Supplementary Figure 1D, F, I-J**). RD was the most sensitive metric for the left AF. RD and FA^TDF^ were similarly sensitive for left ILF (**Supplementary Figure 1G-H**).

**Figure 5.**
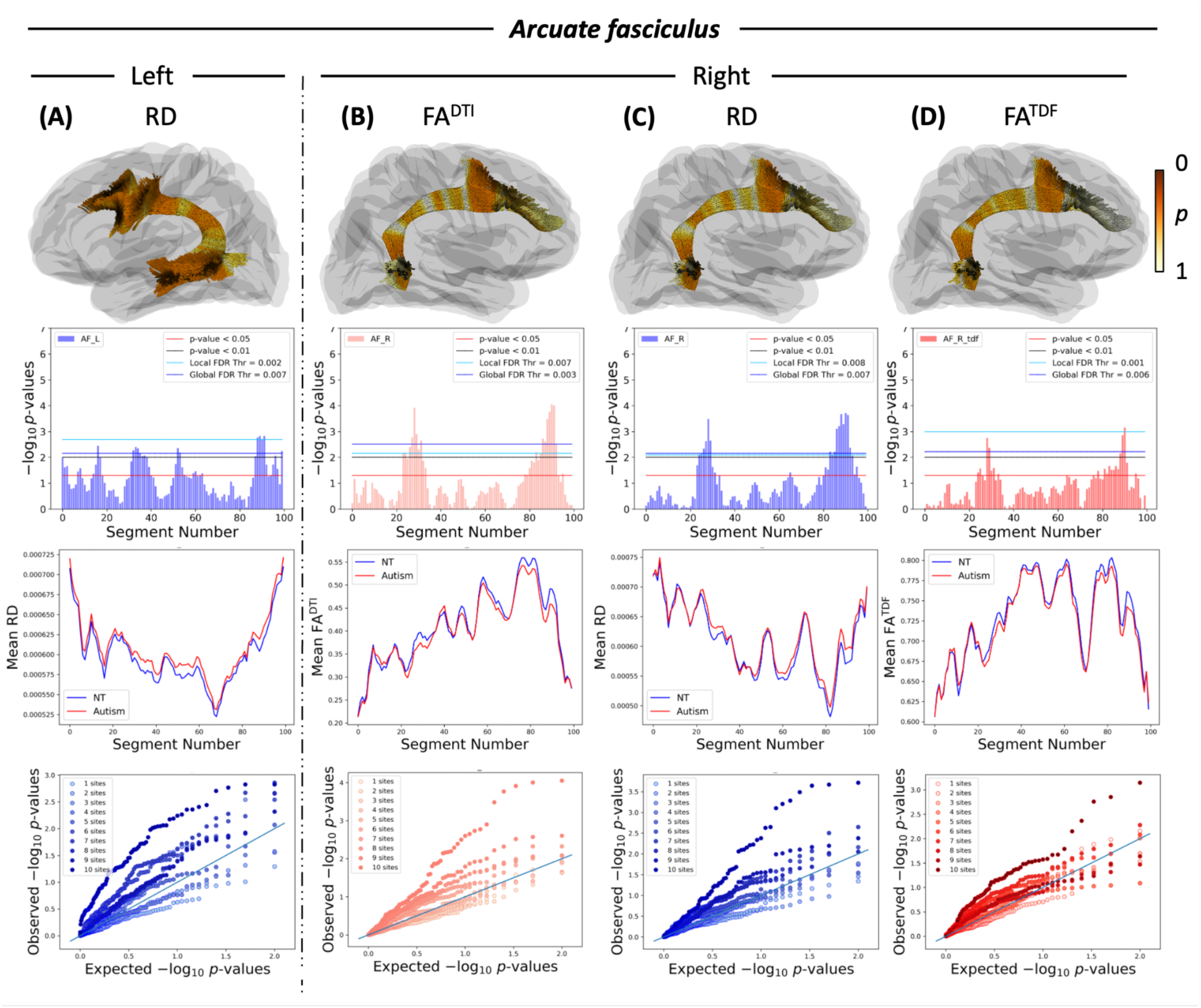
White matter microstructural alterations in the arcuate fasciculus (AF) in autism compared to neurotypical controls. Each column represents a different white matter metric. (A) RD for the left AF, (B) FA^DTI^ for the right AF, (C) RD for the right AF, and (D) FA^TDF^ for the right AF. The first row is a 3D representation of the AF illustrating the anatomical location and corresponding *p*-values of the corresponding microstructural metric; the colorbar shows the *p*-values, where darker orange is a lower *p*-value nearing 0 and lighter yellow is a *p*-value of 1. The second row depicts the negative logarithms of *p*-values for each segment along the tract when contrasting the autism and neurotypical groups; the superior portion of the tract is segment 1, and the segment number increases along the length of the tract up to segment 100 on the inferior portion of the tract. The third row depicts mean plots showing the mean microstructural metric for each group within each segment with autism in red and neurotypicals in blue. The fourth row consists of QQ plots that display the effect of adding more cohorts, where including additional cohorts increases sensitivity to white matter differences. FA^DTI^, fractional anisotropy; RD, radial diffusivity; FA^TDF^, tensor distribution function fractional anisotropy.

**Figure 6.**
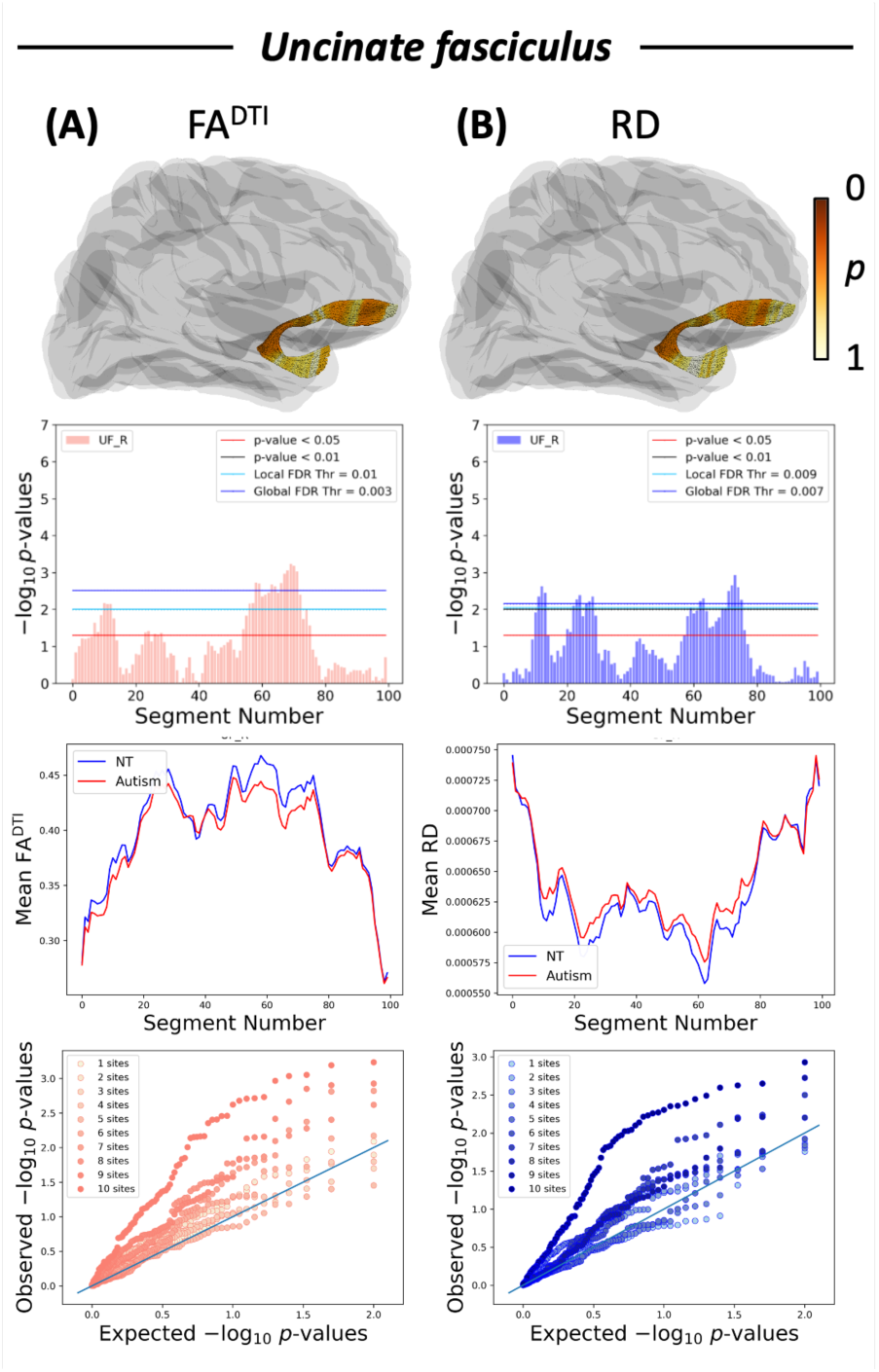
White matter microstructural alterations in the uncinate fasciculus (UF) in autism compared to neurotypical controls. Each column represents a different white matter metric (A) FA^DTI^ and (B) RD. The first row is a 3D representation of the UF illustrating the anatomical location and corresponding *p*-values of the corresponding microstructural metric. The colorbar shows the *p*-values, where darker orange is a lower *p*-value nearing 0 and lighter yellow is a *p*-value of 1. The second row depicts the negative logarithms of *p*-values for each segment along the tract when contrasting the autism and neurotypical groups; the anterior superior portion of the tract is segment 1, and the segment number increases along the length of the tract up to segment 100 on the anterior inferior portion of the tract. The third row depicts mean plots showing the mean microstructural metric for each group within each segment with autism in red and neurotypicals in blue. The fourth row consists of QQ plots that display the effect of adding more cohorts, where including additional cohorts increases sensitivity to white matter differences. FA^DTI^, fractional anisotropy; RD, radial diffusivity

**Figure 7.**
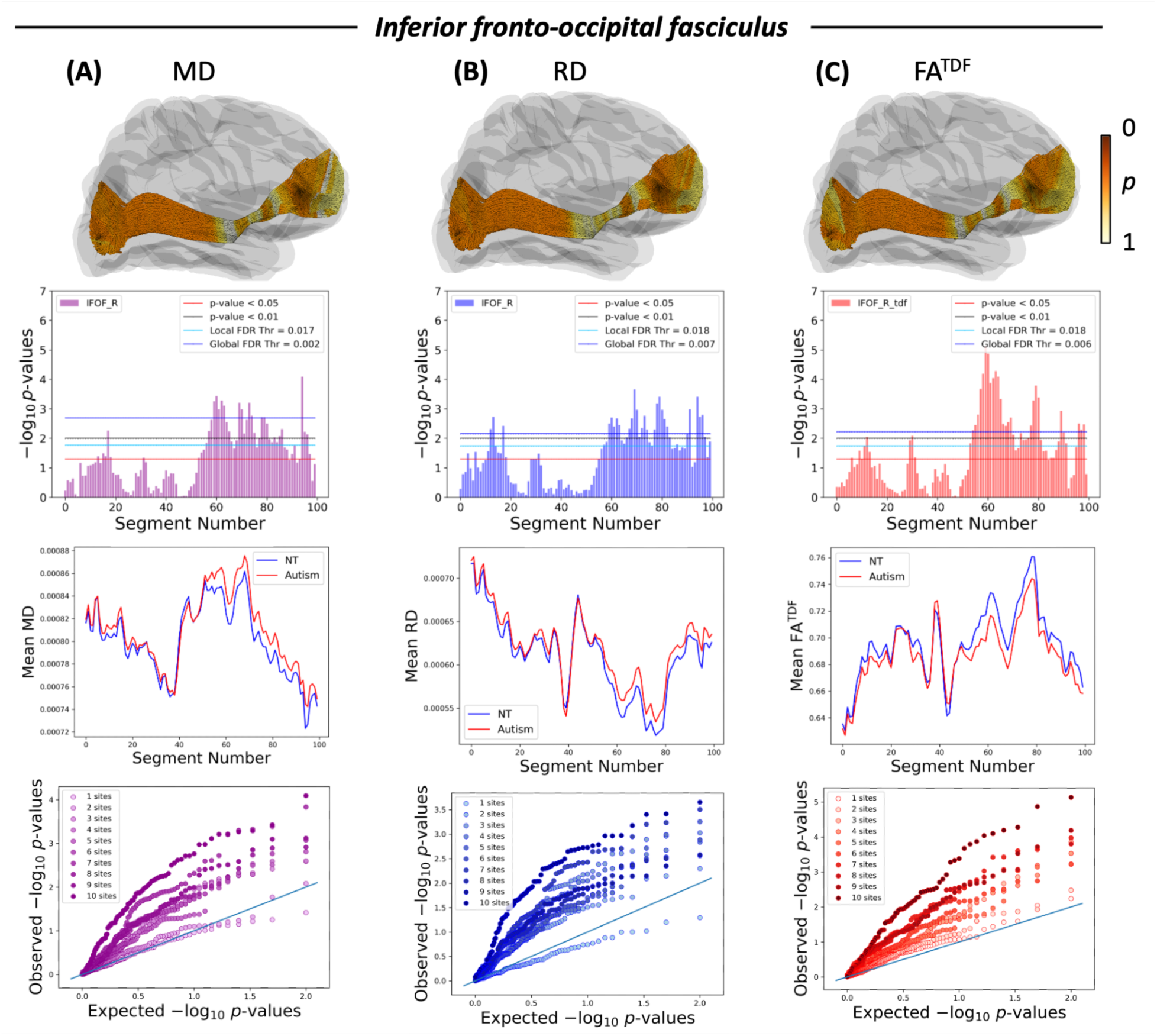
White matter microstructural alterations in the inferior fronto-occipital fasciculus (IFOF) in autism compared to neurotypical controls. Each column represents a different white matter metric (A) MD, (B) RD, and (C) FA^TDF^. The first row is a 3D representation of the IFOF illustrating the anatomical location and corresponding *p*-values of the corresponding microstructural metric; the colorbar shows the *p*-values, where darker orange is a lower *p*-value nearing 0 and lighter yellow is a *p*-value of 1. The second row depicts the negative logarithms of *p*-values for each segment along the tract when contrasting the autism and neurotypical groups; the anterior portion of the tract is segment 1, and the segment number increases along the length of the tract up to segment 100 on the posterior portion of the tract. The third row depicts mean plots showing the mean microstructural metric for each group within each segment with autism in red and neurotypicals in blue.The fourth row consists of QQ plots that display the effect of adding more cohorts, where including additional cohorts increases sensitivity to white matter differences. MD, mean diffusivity; RD, radial diffusivity; FA^TDF^, tensor distribution function fractional anisotropy

**Figure 8.**
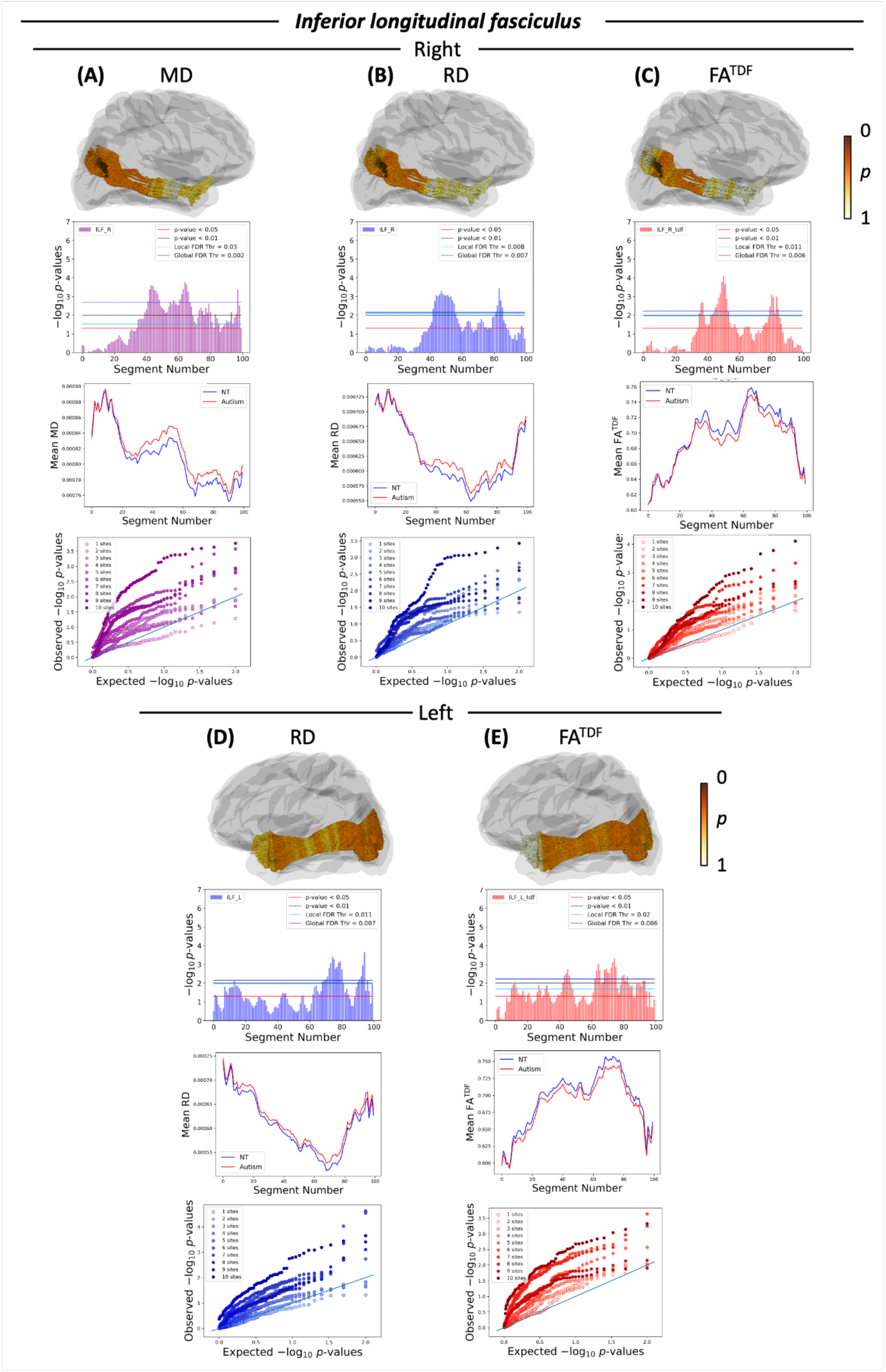
White matter microstructural alterations in the inferior longitudinal fasciculus (ILF) in autism compared to neurotypical controls. Each column represents a different white matter metric (A) MD, (B) RD, and (C) FA^TDF^ for the right ILF and (D) RD and (E) FA^TDF^ for the left ILF. The first row is a 3D representation of the ILF illustrating the anatomical location and corresponding *p*-values of the corresponding microstructural metric; the colorbar shows the *p*-values, where darker orange is a lower *p*-value nearing 0 and lighter yellow is a *p*-value of 1. The second row depicts the negative logarithms of *p*-values for each segment along the tract when contrasting the autism and neurotypical groups; the anterior portion of the tract is segment 1, and the segment number increases along the length of the tract up to segment 100 on the posterior portion of the tract. The third row depicts mean plots showing the mean microstructural metric for each group within each segment with autism in red and neurotypicals in blue. The fourth row consists of QQ plots that display the effect of adding more cohorts, where including additional cohorts increases sensitivity to white matter differences. MD, mean diffusivity; RD, radial diffusivity; FA^TDF^, tensor distribution function fractional anisotropy.

**Figure 9.**
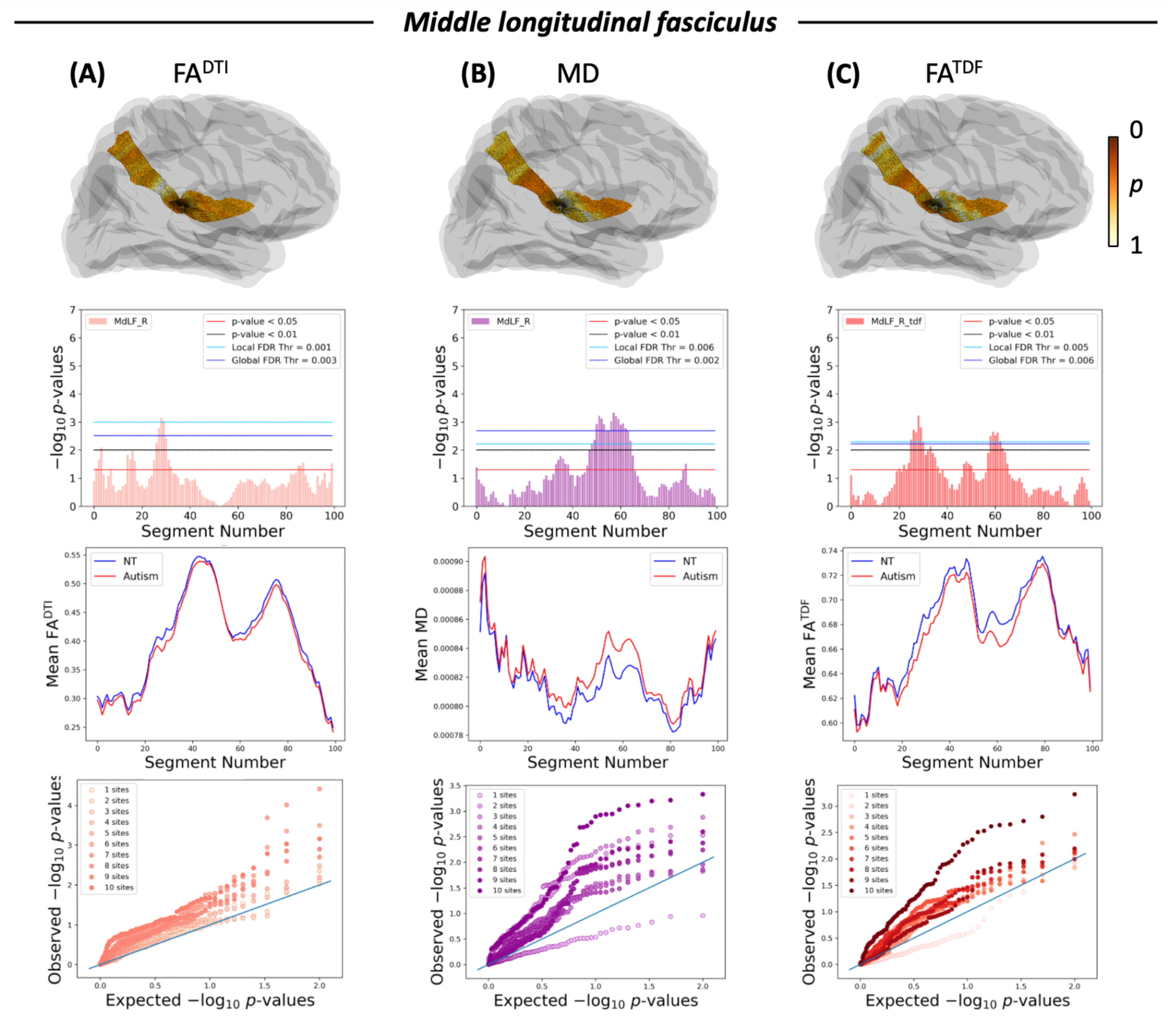
White matter microstructural alterations in the middle longitudinal fasciculus (MdLF) in autism compared to neurotypical controls. Each column represents a different white matter metric (A) FA^DTI^, (B) MD, and (C) FA^TDF^. The first row is a 3D representation of the MdLF illustrating the anatomical location and corresponding *p*-values of the corresponding microstructural metric; the colorbar shows the *p*-values, where darker orange is a lower *p*-value nearing 0 and lighter yellow is a *p*-value of 1. The second row depicts the negative logarithms of *p*-values for each segment along the tract when contrasting the autism and neurotypical groups; the anterior inferior portion of the tract is segment 1, and the segment number increases along the length of the tract up to segment 100 on the posterior superior portion of the tract. The third row depicts mean plots showing the mean microstructural metric for each group within each segment with autism in red and neurotypicals in blue. The fourth row consists of QQ plots that display the effect of adding more cohorts, where including additional cohorts increases sensitivity to white matter differences. FA^DTI^, fractional anisotropy, MD, mean diffusivity; FA^TDF^, tensor distribution function fractional anisotropy

In supplemental analyses, we tested the robustness of our results by completing all analyses when statistically adjusting for head motion. As a whole, the implicated white matter tracts and metrics remained nearly identical. All commissural tracts continued to exhibit significant alterations in autism, with the same microstructure metrics exhibiting significant differences as in our primary analyses. Association tracts similarly continued to display significant alterations that were consistent with the metrics found in our primary analyses, with the exception of FA^DTI^ for the right MdLF which no longer attained statistical significance (**Supplementary Figure 2-9**).

## 4. DISCUSSION

We finely mapped white matter microstructure alterations in one of the largest autism tractometry samples to date using the refined along-tract analysis framework, BUAN, and the advanced TDF model. All three commissural tracts exhibited spatially-specific reductions in FA – commonly interpreted as white matter integrity – in autism, as well as localized elevations in diffusivity within each commissural tract. Association tracts similarly had lower white matter FA and higher diffusivity in autism, with these microstructure alterations observed in localized segments of nearly every association tract examined. These results highlight the complexity of white matter alterations in autism by demonstrating that such alterations are localized within individual tracts but not confined to a specific type of tract or region of the brain.

Our finding that autism was associated with reduced FA^DTI^ and elevated MD compared to neurotypical controls, on average, is in line with prior research (Travers et al. 2012; Di et al. 2018; Zhao et al. 2022). As FA^DTI^ is associated with intact cell membranes, among other underlying biological processes, lower FA^DTI^ in autism may indicate lower microstructural integrity, a reduced coherence of fibers, more crossing fibers, and/or altered neuroinflammation (Travers et al. 2012; Soares et al. 2013; Lord et al. 2020; Hirota and King 2023). Higher MD and RD in the autism group suggest greater white matter water diffusion in autism, which may result from reduced or disrupted myelination, increased fiber spacing, thicker axon diameters, and/or less dense axons (Song et al. 2005; Travers et al. 2012; Soares et al. 2013). Taken together, these DTI results suggest that autism may be associated with inefficient neuronal communication. Although DTI provides valuable insights into white matter microstructure in the brain, its inability to resolve complex fiber configurations results in simplified biological interpretations that do not fully capture connectivity patterns (Basser et al. 1994; Travers et al. 2012; Soares et al. 2013). To address the limitations of the DTI model, we also used the TDF-derived measure, FA^TDF^ (Leow et al. 2009; Nir et al. 2017). FA^TDF^ was also lower in autism than in neurotypical controls, on average, further supporting the presence of reduced white matter integrity in autism. Additionally, FA^TDF^ demonstrated more sensitivity in distinguishing between the autism and neurotypical groups for the commissural tracts and some association tracts, compared to FA^DTI^. This enhanced sensitivity in some tracts aligns with findings from other studies where FA^TDF^ was more sensitive in detecting subtle neural differences (Nir et al. 2017; Lawrence et al. 2021; Benavidez et al. 2024). In sum, the observed pattern of microstructural alterations when using both the DTI and TDF models suggests reduced interhemispheric and intrahemispheric communication in autism (Travers et al. 2012; Soares et al. 2013).

Our finding of altered commissural tract microstructure aligns with prior research indicating that the corpus callosum is affected in autism (Travers et al. 2012; Aoki et al. 2013; Di et al. 2018; Kilroy et al. 2022; Weber et al. 2022; Minnigulova et al. 2023). Using our along-tract mapping methods, we were able to more precisely localize these differences to the lateral regions of the *forceps major* and the medial regions of the *forceps minor* and midbody of the corpus callosum. The corpus callosum is the largest commissural tract, connecting the left and right hemispheres, and plays a critical role in integrating and transferring information between hemispheres (Catani and Schotten 2012; Sakai et al. 2017; Goldstein et al. 2024). The *forceps major* connects the left and right occipital lobes and is involved in visual processing (Catani and Schotten 2012; Ronderos et al. 2024). The *forceps minor* connects the prefrontal regions across hemispheres and is linked to executive function (Catani and Schotten 2012; Mamiya et al. 2018; Costanzo et al. 2022). The midbody of the corpus callosum connects the parietal lobes and premotor and precentral cortex and is associated with sensorimotor and cognitive functions (Hofer and Frahm 2006; Machado et al. 2007; Paul 2011; Catani and Schotten 2012; Mathew et al. 2013). Given the distinct roles of these commissural tracts, the commissural microstructural alterations found in this study may be related to a range of behavioral characteristics observed in autism, including sensory processing differences, executive function challenges, and socioemotional processing challenges (Mathew et al. 2013; Craig et al. 2016; Demetriou et al. 2018; Mamiya et al. 2018; Costanzo et al. 2022; Ronderos et al. 2024).

Our association tract results are in line with prior autism studies suggesting altered white matter microstructure in such tracts (Travers et al. 2012; Aoki et al. 2013; Zhao et al. 2022). Association tracts connect different cortical regions within the same hemisphere and are critical for a range of functions, including supporting higher-order cognitive functions, integrating sensory information, and coordinating complex behaviors (Weiller et al. 2021). The AF connects posterior temporal and frontal regions and is associated with language processing (Yeh et al. 2013; Barbeau et al. 2020); the AF is notably lateralized, such that in most individuals the left AF plays a key role in language production and comprehension, whereas the right AF supports broader language processing (Lebel and Beaulieu 2009; Barbeau et al. 2020). The UF connects the anterior temporal lobe to the lateral orbitofrontal cortex and frontal pole, passing through the insula, and is important for processing socioemotional information (Von Der Heide et al. 2013). Therefore, the microstructural alterations we found in the AF and UF may be associated with the social communication and socioemotional differences in autism (Von Der Heide et al. 2013; Robertson and Baron-Cohen 2017; Barbeau et al. 2020; American Psychiatric Association 2022; Hirota and King 2023). The ILF connects the occipital lobe to the anterior temporal lobe and is primarily involved in visual processing, including supporting a wide range of cognitive and affective processes related to vision (Herbet et al. 2018; Zemmoura et al. 2021). The IFOF connects the occipital lobe to the inferior frontal lobe and is involved in semantic processing (Martino et al. 2010; Conner et al. 2018). The MdLF is a tract that connects the superior temporal gyrus with the superior parietal lobe, parieto-occipital lobe, and the angular gyrus (Wang et al. 2013); the MdLF may play a role in processing and learning auditory information (Latini et al. 2021). The microstructural differences that we observed in the ILF, IFOF, and MdLF tracts may thus relate to altered sensory and semantic processing in autism (Martino et al. 2010; Conner et al. 2018; Herbet et al. 2018; Lord et al. 2020; Key and D’Ambrose Slaboch 2021; Latini et al. 2021; Zemmoura et al. 2021). Overall, alterations in the microstructure of association tracts may disrupt the integration of sensorimotor and socioemotional information, contributing to core behavioral features of autism (Joseph et al. 2014; Li et al. 2017; Barbeau et al. 2020; Olivé et al. 2022; Pitzianti et al. 2023). As our association tract findings were predominantly in the right hemisphere, our results may build on previous brain lateralization morphometry studies in autism that have identified atypical lateralization, suggesting that neurodevelopmental differences associated with brain morphometric asymmetry may propagate through disruptions in structural connectivity (Postema et al. 2019; Floris et al. 2021; Sha et al. 2022).

Taken together, our results expand on previous structural and functional connectivity studies to support theories that autism is characterized by altered brain connectivity (Picci et al. 2016; Vasa et al. 2016; Ilioska et al. 2023; Moreau et al. 2023). As we observed connectivity differences in multiple pathways distributed across the brain, these results indicate potential neurodevelopmental white matter alterations that reflect whole-brain differences in white matter microstructure (Von Der Heide et al. 2013; Picci et al. 2016; Barbeau et al. 2020). Our finding of widespread commissural and association tract differences in autism also align with and expand on these theories by suggesting that altered connectivity between brain regions may contribute to a myriad of the behavioral characteristics associated with autism, including differences in social communication, sensory processing, executive function, and socioemotional functioning. As tracts related to both social and sensory processing are implicated by our analyses, these results furthermore lend support to both social and sensorimotor theories of autism (Hannant et al. 2016; Robertson and Baron-Cohen 2017).

The present study has several strengths. The advanced along-tract analysis method, BUAN, allows for a more precise evaluation of white matter tract segments by not assuming that the entire tract exhibits the same pattern of differences (Chandio et al. 2020). Our results using BUAN therefore provide an important expansion on prior autism studies which reported averaged white matter microstructure differences. Moreover, the refined TDF model used here addresses the limitations of the DTI model by more accurately capturing complex fiber orientations (Leow et al. 2009; Nir et al. 2017). Our use of the advanced single-shell model, TDF, also allows for compatibility with future studies leveraging large-scale single-shell archival datasets and newly collected single-shell datasets in complex study populations that may benefit from short scan acquisition times. The current study is also, to the best of our knowledge, the largest tractography study to date examining individual tracts in autism (Hrdlicka et al. 2019; Bassell et al. 2020; Kilroy et al. 2022; Minnigulova et al. 2023); our large sample size importantly enhances the reliability and robustness of the reported results (Button et al. 2013; Marek et al. 2022). Finally, the analysis of both commissural and association tracts across the brain provide a more comprehensive understanding of white matter in autism by revealing that white matter alterations in autism extend to include multiple different types of tracts throughout the brain. Future studies should collect multi-shell data to examine additional microstructure metrics, such as those provided by the neurite orientation dispersion and density imaging (NODDI) model, to provide further insights into white matter differences in autism (Zhang et al. 2012; Carper et al. 2017; Andica et al. 2021). Such future work should also extend these findings by examining additional white matter tracts, including limbic, projection, and cerebellar tracts, as well as additional association tracts not included in the originally reported BUAN pipeline (Supekar et al. 2018; Chandio et al. 2020). The potential effects of age, sex, and autism symptom severity on white matter microstructure in autism also warrant further investigation, as well as pubertal effects during development on white matter in autism (Herting et al. 2017; Lebel and Deoni 2018; Lawrence et al. 2019; Lei et al. 2019; Lawrence et al. 2020; Andrews et al. 2021; Piekarski et al. 2023; Villalón-Reina et al. 2023). As part of this future research, detailed behavioral measures that align with the functions of the altered tracts should be collected and analyzed, including measures of sensory processing, executive function, language, and socioemotional processing. Lastly, recruiting participants from a wider range of backgrounds will be important for ensuring that the observed findings are representative of the global population (Garcini et al. 2022; Goldfarb and Brown 2022).

In summary, our findings in a large sample of 365 autistic and neurotypical participants indicate that autism is associated with significant white matter alterations across commissural and association tracts. Importantly, these alterations are localized within specific segments of these tracts rather than uniformly affecting the entire tract. The advanced dMRI methods BUAN and TDF rigorously captured these microstructural differences in a spatially precise and sensitive manner, demonstrating the promise of these methods for advancing future research in autism and other brain-based conditions. As a whole, our study reveals nuanced and widespread microstructural alterations in autism that significantly impact white matter tracts across the brain.

## 5. ACKNOWLEDGMENTS

This research was supported by the National Institutes of Health (award K01MH135160 to K.E.L. and RF1AG057892 to P.M.T.), the Momental Foundation (Mistletoe Research Fellowship to K.E.L.), and the Asan Foundation (Biomedical Science Scholarship to G.S.K.). Research reported in this publication was also supported by the Office Of The Director, of the National Institutes of Health under Award Number S10OD032285. The authors have no relevant financial or non-financial interests to disclose.

Data used in the preparation of this article reside in part in the NIH-supported NIMH Data Repositories, a collaborative informatics system created by the National Institutes of Health to provide a national resource to support and accelerate research in mental health related conditions. Dataset identifiers: nda2021 (6 sites/scanners) and nda1906 (1 site/scanner). We thank the ABIDE II team for providing the data (at http://fcon_1000.projects.nitrc.org/indi/abide). Dataset identifiers: Trinity College Dublin (1 site/scanner), San Diego State University (1 site/scanner), Barrow Neurological Institute (1 site/scanner). This manuscript reflects the views of the authors and does not reflect the opinions or views of the NIH or of the Submitters submitting original data to the NIMH Data Repositories or ABIDE II.

