## Supplementary for "Mapping Along-Tract White Matter Microstructural Differences in Autism"

### **Supplementary Figures/Tables**

**Supplementary Table 1**. Participant demographics by site

**Supplementary Table 2**. Scanner information by site

**Supplementary Figure 1**. Quantile-Quantile plot of each microstructural metric for each significant white matter tract

**Supplementary Figure 2**. White matter microstructural alterations in the *forceps minor* in autism compared to neurotypical controls accounting for motion

**Supplementary Figure 3**. White matter microstructural alterations in the midbody of corpus callosum in autism compared to neurotypical controls accounting for motion

**Supplementary Figure 4**. White matter microstructural alterations in the *forceps major* in autism compared to neurotypical controls accounting for motion

**Supplementary Figure 5**. White matter microstructural alterations in the arcuate fasciculus in autism compared to neurotypical controls accounting for motion

**Supplementary Figure 6**. White matter microstructural alterations in the uncinate fasciculus in autism compared to neurotypical controls accounting for motion

**Supplementary Figure 7**. White matter microstructural alterations in the inferior fronto-occipital fasciculus in autism compared to neurotypical controls accounting for motion

**Supplementary Figure 8**. White matter microstructural alterations in the inferior longitudinal fasciculus in autism compared to neurotypical controls accounting for motion

**Supplementary Figure 9**. White matter microstructural alterations in the middle longitudinal fasciculus in autism compared to neurotypical controls accounting for motion

#### **Supplementary Table 1**. Participant demographics by site

|  | **Total**  **(N=365)** | **BNI**  **(N=19)** | **SDSU**  **(N=50)** | **TCD**  **(N=37)** | **NDA1906**  **(N=25)** | **NDA2021-HT**  **(N=38)** | **NDA2021-SP**  **(N=43)** | **NDA2021-ST**  **(N=21)** | **NDA2021-UP**  **(N=23)** | **NDA2021-UT**  **(N=60)** | **NDA2021-YT**  **(N=49)** |
| --- | --- | --- | --- | --- | --- | --- | --- | --- | --- | --- | --- |
| **Diagnosis** | |  |  |  |  |  |  |  |  |  |  |
| Autism | 195 (53 %) | 11 (58 %) | 28 (56 %) | 17 (46 %) | 18 (72 %) | 23 (61 %) | 19 (44 %) | 15 (71 %) | 8 (35 %) | 31 (52 %) | 25 (51 %) |
| Neurotypical | 170 (47 %) | 8 (42 %) | 22 (44 %) | 20 (54 %) | 7 (28 %) | 15 (39 %) | 24 (56 %) | 6 (29 %) | 15 (65 %) | 29 (48 %) | 24 (49 %) |
| **Sex** | |  |  |  |  |  |  |  |  |  |  |
| F | 124 (34 %) | 0 (0 %) | 8 (16 %) | 0 (0 %) | 0 (0 %) | 15 (39 %) | 21 (49 %) | 12 (57 %) | 12 (52 %) | 26 (43 %) | 30 (61 %) |
| M | 241 (66 %) | 19 (100 %) | 42 (84 %) | 37 (100 %) | 25 (100 %) | 23 (61 %) | 22 (51 %) | 9 (43 %) | 11 (48 %) | 34 (57 %) | 19 (39 %) |
| **Age (years)** | |  |  |  |  |  |  |  |  |  |  |
| Mean (SD) | 14 (± 3.5) | 20 (± 1.7) | 13 (± 3.1) | 15 (± 3.2) | 12 (± 5.1) | 12 (± 3.1) | 13 (± 2.5) | 12 (± 2.9) | 13 (± 3.3) | 13 (± 3.1) | 14 (± 2.6) |
| **Mean Relative Motion (mm)** | |  |  |  |  |  |  |  |  |  |  |
| Mean (SD) | 0.29  (± 0.14) | 0.23  (± 0.051) | 0.26  (± 0.059) | 0.31  (± 0.12) | 0.27  (± 0.060) | 0.27  (± 0.11) | 0.34  (± 0.17) | 0.30  (± 0.16) | 0.23  (± 0.13) | 0.33  (± 0.16) | 0.32  (± 0.16) |
| **FSIQ** | |  |  |  |  |  |  |  |  |  |  |
| Mean (SD) | 110 (± 18) | 110 (± 14) | 100 (± 14) | 120 (± 14) | 94 (± 16) | 100 (± 19) | 110 (± 22) | 110 (± 20) | 110 (± 17) | 100 (± 17) | 100 (± 15) |
| **SRS T Score**^a^ | |  |  |  |  |  |  |  |  |  |  |
| Mean (SD) | 61 (± 18) | 64 (± 17) | 65 (± 22) | 60 (± 20) | 67 (± 19) | 60 (± 16) | 57 (± 18) | 67 (± 18) | 54 (± 17) | 60 (± 17) | 59 (± 17) |
| **ADOS CSS**^a^ | |  |  |  |  |  |  |  |  |  |  |
| Mean (SD) | 12 (± 5.1) | NA | NA | NA | 15 (± 7.9) | 11 (± 4.6) | 13 (± 4.5) | 11 (± 3.6) | 13 (± 3.5) | 12 (± 4.0) | 11 (± 5.1) |

BNI, SDSU, TCD are from the ABIDE-II dataset. NDA1906, NDA2021-HT, NDA2021-SP, NDA2021-ST, NDA2021-UP, NDA2021-UT, NDA2021-YT are from the NIMH Data Archive. BNI, Barrow Neurological Institute; SDSU, San Diego State University; TCD, Trinity College Dublin; HT, Harvard Trio; SP, Seattle Prisma; ST, Seattle Trio; UP, UCLA Prisma; UT, UCLA Trio; YT, Yale Trio; F, Female; M, Male; SD, standard deviation; FSIQ, Full Scale Intelligence Quotient; SRS, Social Responsiveness Scale; ADOS CSS, Autism Diagnostic Observation Schedule Calibrated Severity Score

^a^The mean value is calculated based only on subjects with available data for this measure.

#### **Supplementary Table 2**. Scanner information by site

|  | **Scanner** | **Field strength**  (T) | **Voxel Size**  (mm) | **b=0 scans** | **Gradient directions** | **B-value**  (mm/s^2^) | **Slices** | **TE**  (ms) | **TR**  (ms) |
| --- | --- | --- | --- | --- | --- | --- | --- | --- | --- |
| **BNI** | Philips | 3 | 1.4x1.4x3.0 | 1 | 32 | 2500 | 48 | 101 | 7850 |
| **SDSU** | GE | 3 | 2.0x2.0x2.0 | 1 | 61 | 1000 | 68 | 81.8 | 8500 |
| **TCD** | Philips | 3 | 2.0x2.0x2.0 | 1 | 61 | 1500 | 65 | 79 | 20244 |
| **NDA1906** | Siemens | 3 | 2.0x2.0x2.5 | 4 | 48 | 1000 | 50 | 91 | 7000 |
| **NDA2021-HT** | Siemens | 3 | 1.98x1.98x2 | 1 | 64 | 1000 | 60 | 93 | 11500 |
| **NDA2021-SP** | Siemens | 3 | 1.98x1.98x2 | 6 | 64 | 1000 | 60 | 74 | 7300 |
| **NDA2021-ST** | Siemens | 3 | 1.98x1.98x2 | 1 | 64 | 1000 | 60 | 93 | 11500 |
| **NDA2021-UP** | Siemens | 3 | 1.98x1.98x2 | 6 | 64 | 1000 | 60 | 74 | 7300 |
| **NDA2021-UT** | Siemens | 3 | 1.98x1.98x2 | 1 | 64 | 1000 | 60 | 93 | 11500 |
| **NDA2021-YT** | Siemens | 3 | 1.98x1.98x2 | 1 | 64 | 1000 | 60 | 93 | 11500 |

BNI, SDSU, TCD are from the ABIDE-II dataset. NDA1906, NDA2021-HT, NDA2021-SP, NDA2021-ST, NDA2021-UP, NDA2021-UT, NDA2021-YT are from the NIMH Data Archive. BNI, Barrow Neurological Institute; SDSU, San Diego State University; TCD, Trinity College Dublin; HT, Harvard Trio; SP, Seattle Prisma; ST, Seattle Trio; UP, UCLA Prisma; UT, UCLA Trio; YT, Yale Trio.

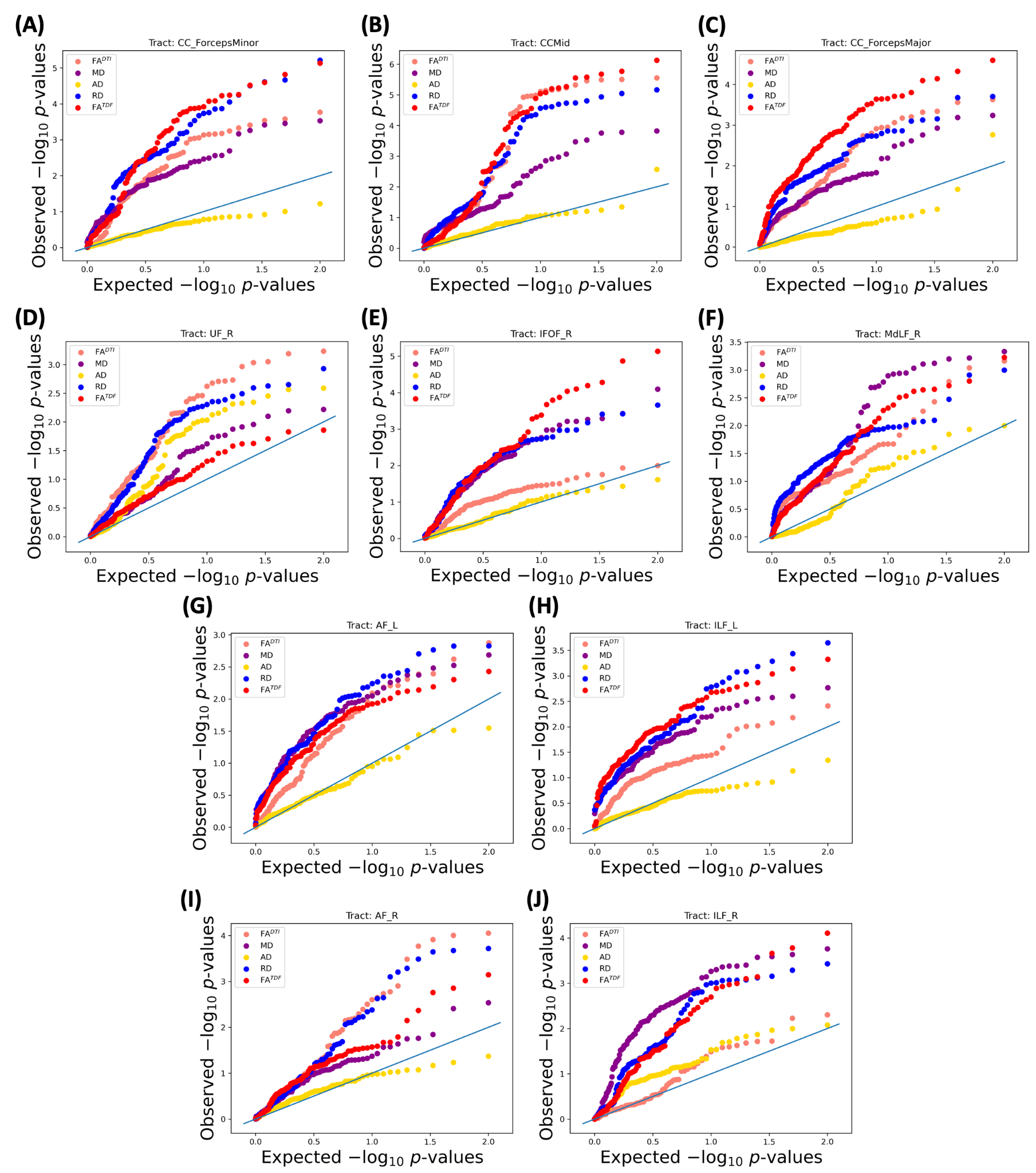

**Supplementary Figure 1**. Quantile-Quantile plot of each microstructural metric for each significant white matter tract. (A) *Forceps minor*, (B) midbody of the corpus callosum, (C) *forceps major*, (D) right uncinate fasciculus, (E) right inferior fronto-occipital fasciculus, (F) right middle longitudinal fasciculus, (G) left arcuate fasciculus, (H) left inferior longitudinal fasciculus, (I) right arcuate fasciculus, and (J) right inferior longitudinal fasciculus. FA^DTI^, fractional anisotropy; MD, mean diffusivity; RD, radial diffusivity; AD, axial diffusivity; FA^TDF^, tensor distribution function fractional anisotropy; AF, arcuate fasciculus; CC, corpus callosum; IFOF, inferior fronto-occipital fasciculus; ILF, inferior longitudinal fasciculus; MdLF, middle longitudinal fasciculus; UF, uncinate fasciculus; L, left; R, right.

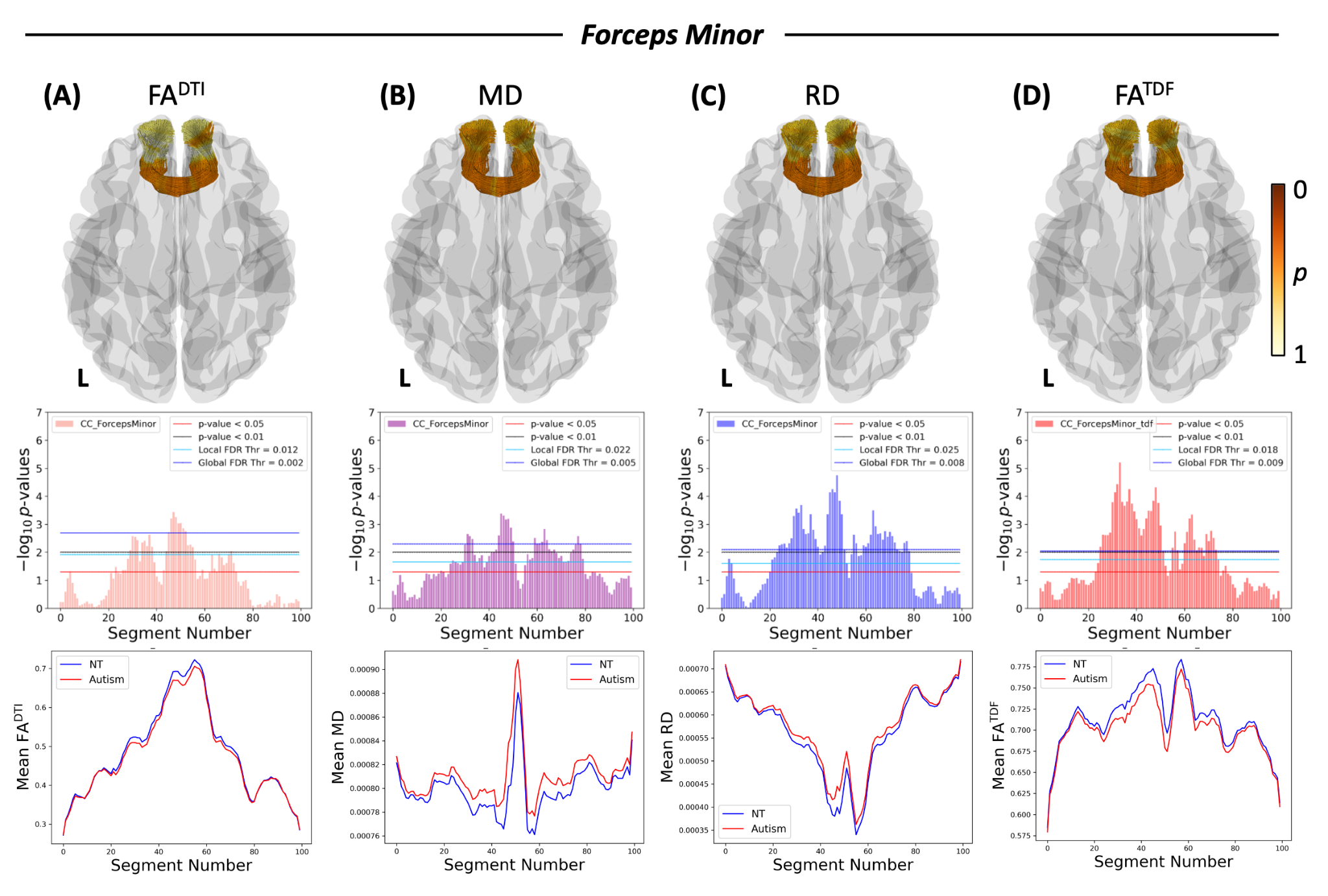

**Supplementary Figure 2.** White matter microstructural alterations in the *forceps minor* in autism compared to neurotypical controls*.* Each column represents a different white matter metric (A) FA^DTI^, (B) MD, (C) RD, and (C) FA^TDF^. The first row is a 3D representation of the *forceps minor* illustrating the anatomical location and corresponding *p*-values of the corresponding microstructural metric; L indicates the left brain hemisphere and the colorbar shows the *p*-values, where darker orange is a lower *p*-value nearing 0 and lighter yellow is a *p*-value of 1. The second row depicts the negative logarithms of *p*-values for each segment along the tract when contrasting the autism and neurotypical groups; the right hemisphere’s anterior portion of the tract is segment 1, and the segment number increases along the length of the tract up to segment 100 on the left hemisphere’s anterior portion of the tract. The third row depicts mean plots showing the mean microstructural metric for each group within each segment with autism in red and neurotypicals in blue. FA^DTI^, fractional anisotropy; MD, mean diffusivity; RD, radial diffusivity; FA^TDF^, tensor distribution function fractional anisotropy

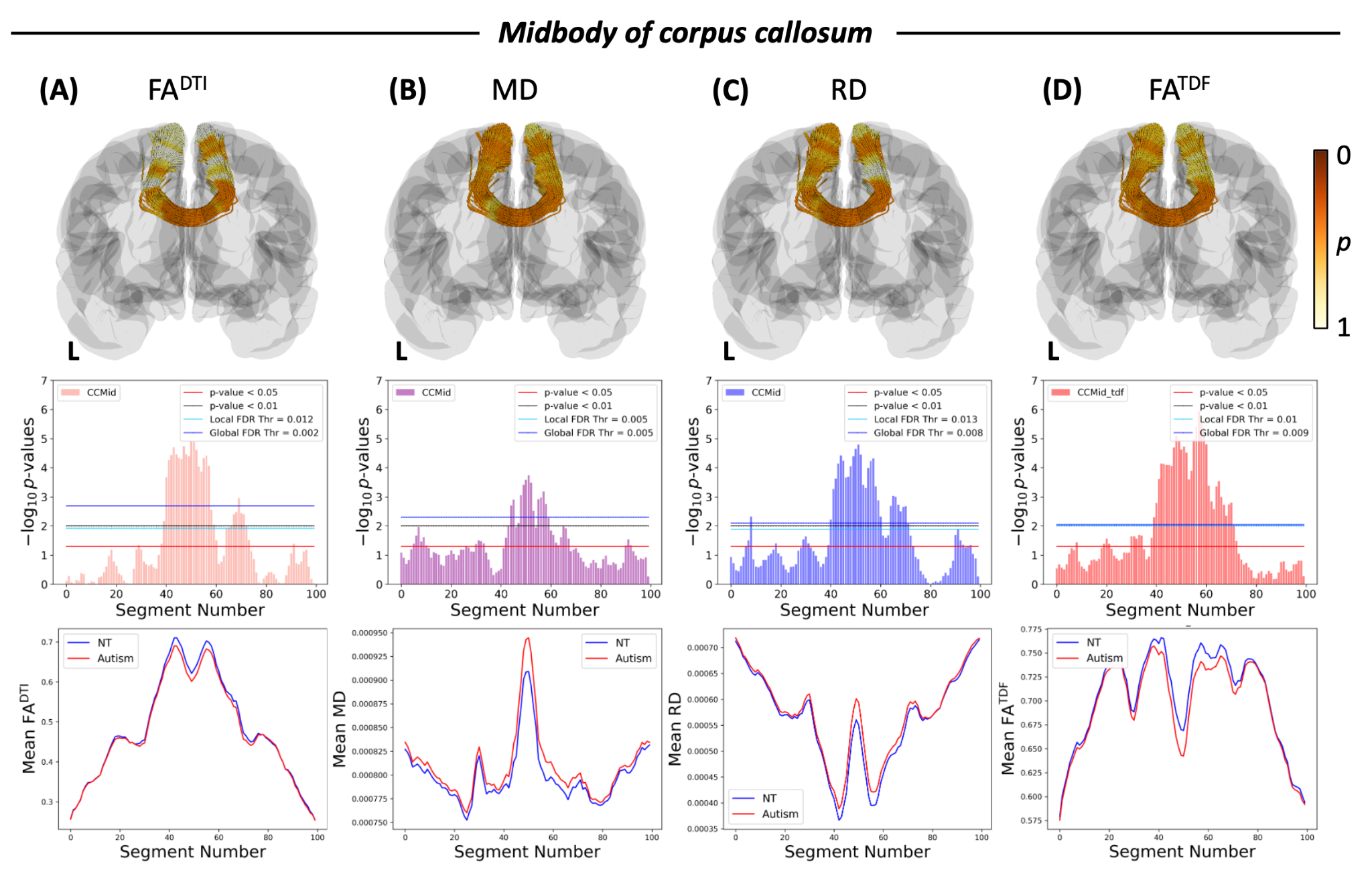

**Supplementary Figure 3.** White matter microstructural alterations in the midbody of the corpus callosum in autism compared to neurotypical controls*.* Each column represents a different white matter metric (A) FA^DTI^, (B) MD, (C) RD, and (C) FA^TDF^. The first row is a 3D representation of the midbody of the corpus callosum illustrating the anatomical location and corresponding *p*-values of the corresponding microstructural metric; L indicates the left brain hemisphere and the colorbar shows the *p*-values, where darker orange is a lower *p*-value nearing 0 and lighter yellow is a *p*-value of 1. The second row depicts the negative logarithms of *p*-values for each segment along the tract when contrasting the autism and neurotypical groups; the left hemisphere’s superior portion of the tract is segment 1, and the segment number increases along the length of the tract up to segment 100 on the right hemisphere’s superior portion of the tract. The third row depicts mean plots showing the mean microstructural metric for each group within each segment with autism in red and neurotypicals in blue. FA^DTI^, fractional anisotropy; MD, mean diffusivity; RD, radial diffusivity; FA^TDF^, tensor distribution function fractional anisotropy

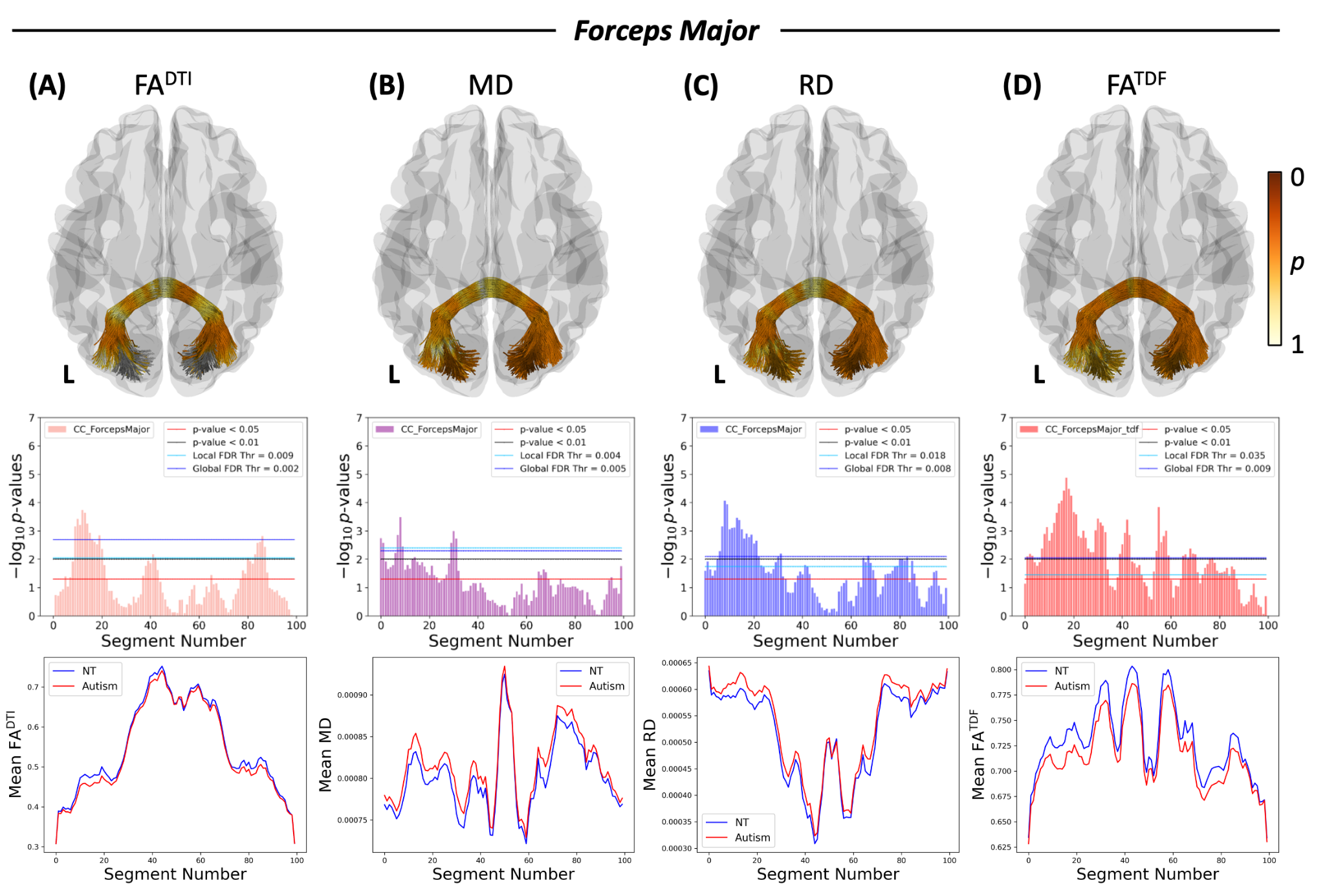

**Supplementary Figure 4.** White matter microstructural alterations in the *forceps major* in autism compared to neurotypical controls*.* Each column represents a different white matter metric (A) FA^DTI^, (B) MD, (C) RD, and (C) FA^TDF^. The first row is a 3D representation of the *forceps major* illustrating the anatomical location and corresponding *p*-values of the corresponding microstructural metric; L indicates the left brain hemisphere and the colorbar shows the *p*-values, where darker orange is a lower *p*-value nearing 0 and lighter yellow is a *p*-value of 1. The second row depicts the negative logarithms of *p*-values for each segment along the tract when contrasting the autism and neurotypical groups; the right hemisphere’s posterior portion of the tract is segment 1, and the segment number increases along the length of the tract up to segment 100 on the left hemisphere’s posterior portion of the tract. The third row depicts mean plots showing the mean microstructural metric for each group within each segment with autism in red and neurotypicals in blue. FA^DTI^, fractional anisotropy; MD, mean diffusivity; RD, radial diffusivity; FA^TDF^, tensor distribution function fractional anisotropy

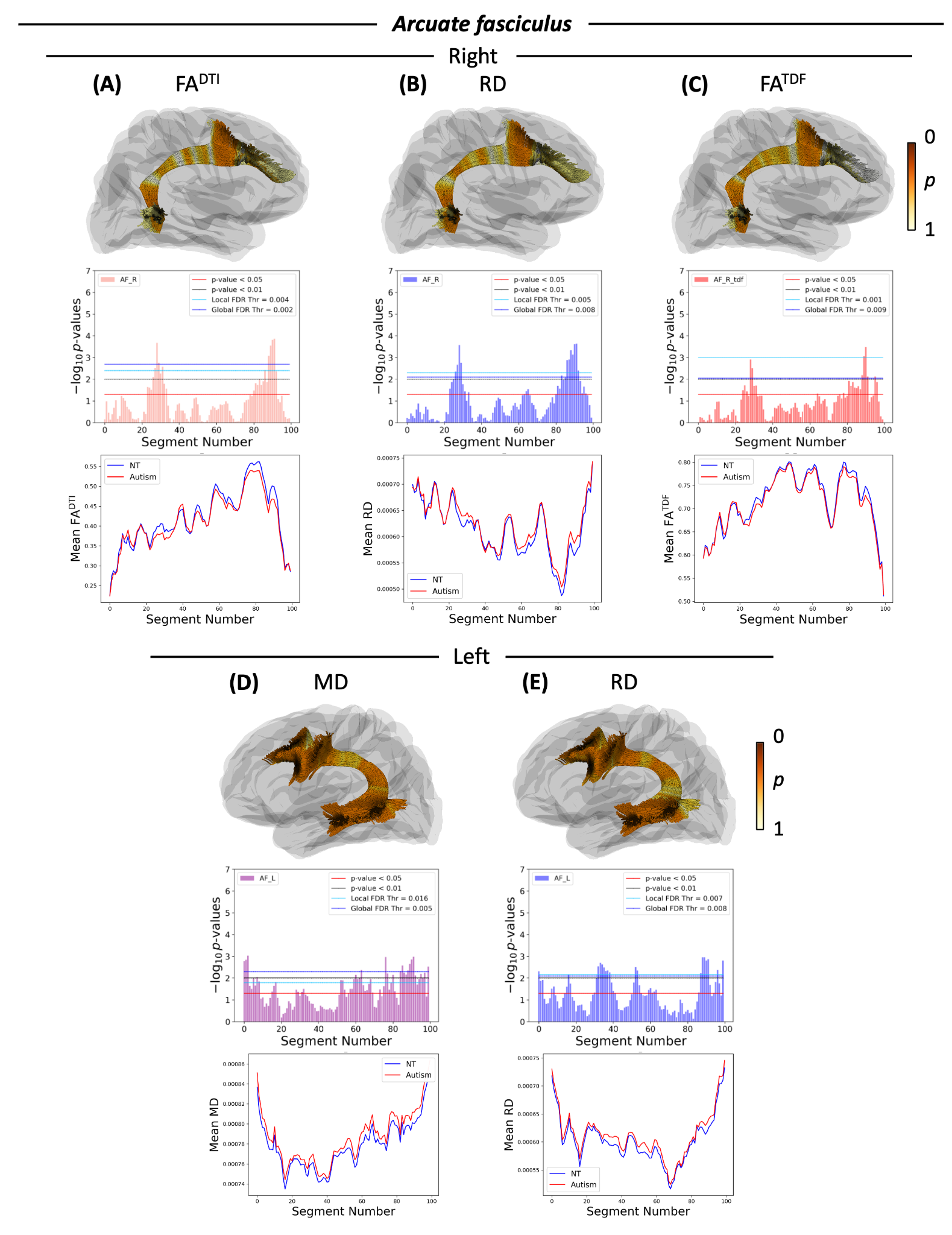

**Supplementary Figure 5.** White matter microstructural alterations in the arcuate fasciculus (AF) in autism compared to neurotypical controls*.* Each column represents a different white matter metric. (A) FA^DTI^ for the right AF, (B) RD for the right AF, (C) FA^TDF^ for the right AF, (D) MD for the left AF, and (E) RD for the left AF. The first row is a 3D representation of the AF illustrating the anatomical location and corresponding *p*-values of the corresponding microstructural metric; the colorbar shows the *p*-values, where darker orange is a lower *p*-value nearing 0 and lighter yellow is a *p*-value of 1. The second row depicts the negative logarithms of *p*-values for each segment along the tract when contrasting the autism and neurotypical groups; the superior portion of the tract is segment 1, and the segment number increases along the length of the tract up to segment 100 on the inferior portion of the tract. The third row depicts mean plots showing the mean microstructural metric for each group within each segment with autism in red and neurotypicals in blue. FA^DTI^, fractional anisotropy; MD, mean diffusivity; RD, radial diffusivity; FA^TDF^, tensor distribution function fractional anisotropy.

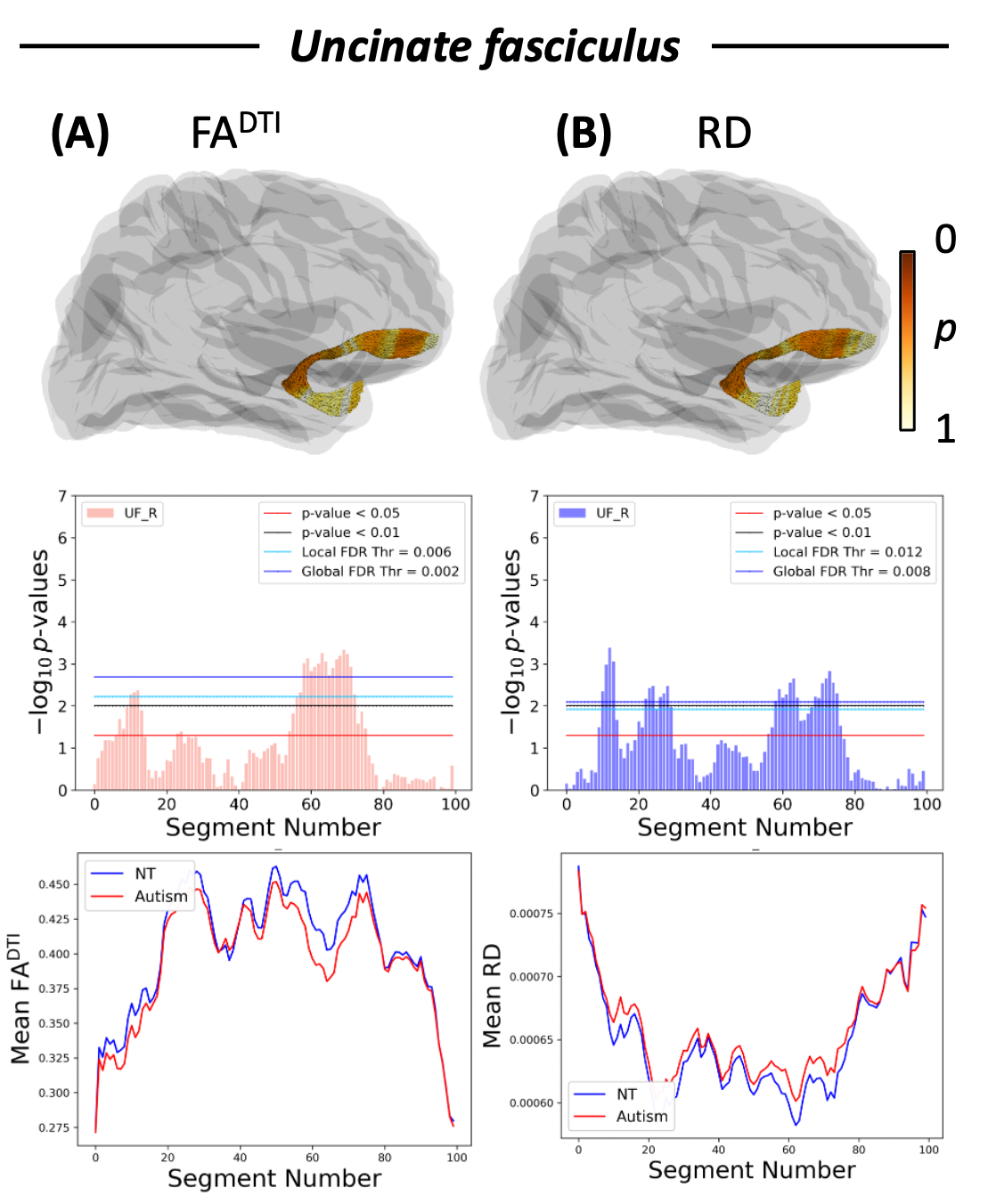

**Supplementary Figure 6.** White matter microstructural alterations in the uncinate fasciculus in autism compared to neurotypical controls*.* Each column represents a different white matter metric (A) FA^DTI^ and (B) RD. The first row is a 3D representation of the uncinate fasciculus illustrating the anatomical location and corresponding *p*-values of the corresponding microstructural metric; the colorbar shows the *p*-values, where darker orange is a lower *p*-value nearing 0 and lighter yellow is a *p*-value of 1. The second row depicts the negative logarithms of *p*-values for each segment along the tract when contrasting the autism and neurotypical groups; the anterior superior portion of the tract is segment 1, and the segment number increases along the length of the tract up to segment 100 on the anterior inferior portion of the tract. The third row depicts mean plots showing the mean microstructural metric for each group within each segment with autism in red and neurotypicals in blue. FA^DTI^, fractional anisotropy; RD, radial diffusivity

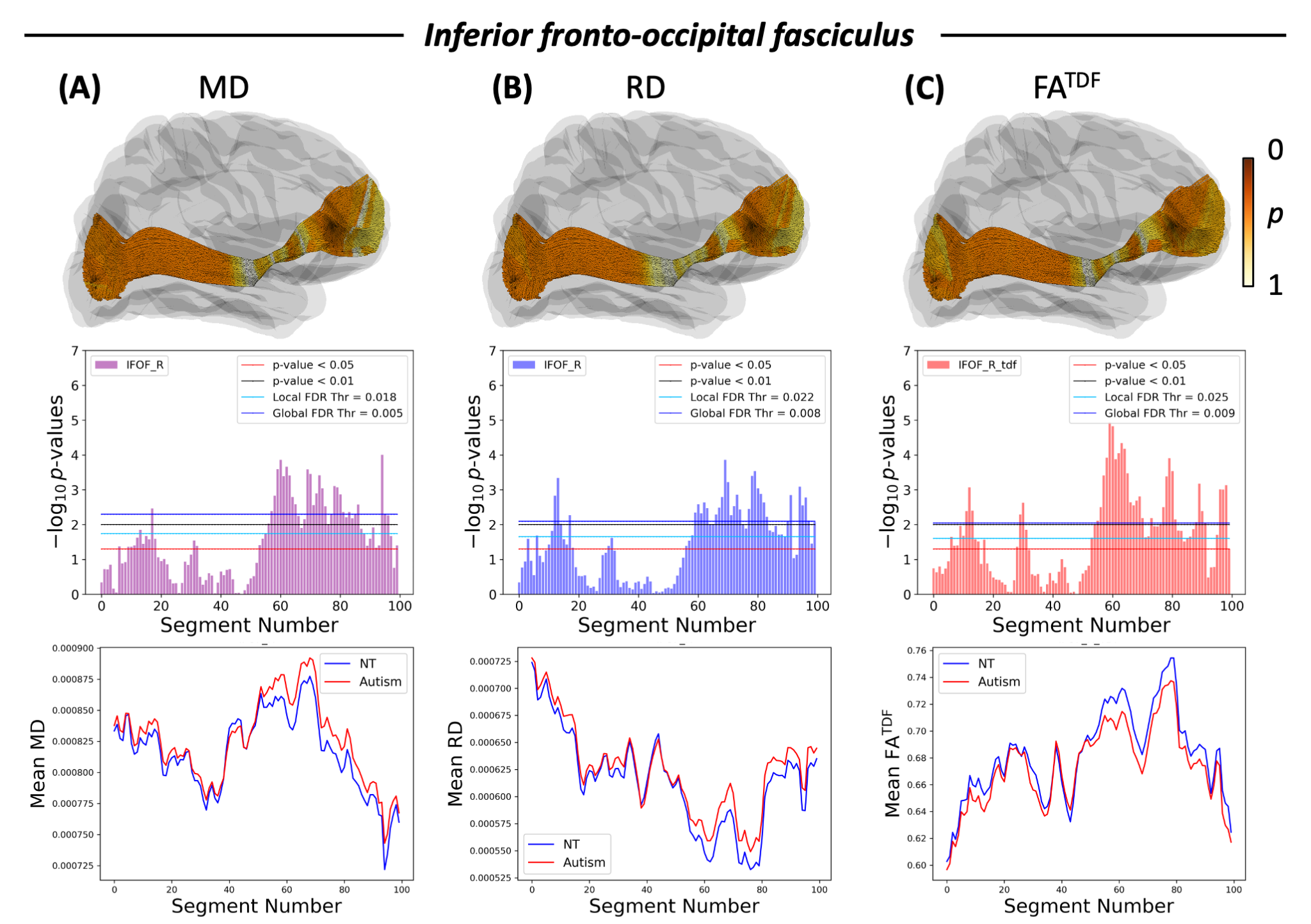

**Supplementary Figure 7.** White matter microstructural alterations in the inferior fronto-occipital fasciculus in autism compared to neurotypical controls*.* Each column represents a different white matter metric (A) MD, (B) RD, and (C) FA^TDF^. The first row is a 3D representation of the inferior fronto-occipital fasciculus illustrating the anatomical location and corresponding *p*-values of the corresponding microstructural metric; the colorbar shows the *p*-values, where darker orange is a lower *p*-value nearing 0 and lighter yellow is a *p*-value of 1. The second row depicts the negative logarithms of *p*-values for each segment along the tract when contrasting the autism and neurotypical groups; the anterior portion of the tract is segment 1, and the segment number increases along the length of the tract up to segment 100 on the posterior portion of the tract. The third row depicts mean plots showing the mean microstructural metric for each group within each segment with autism in red and neurotypicals in blue. MD, mean diffusivity; RD, radial diffusivity; FA^TDF^, tensor distribution function fractional anisotropy

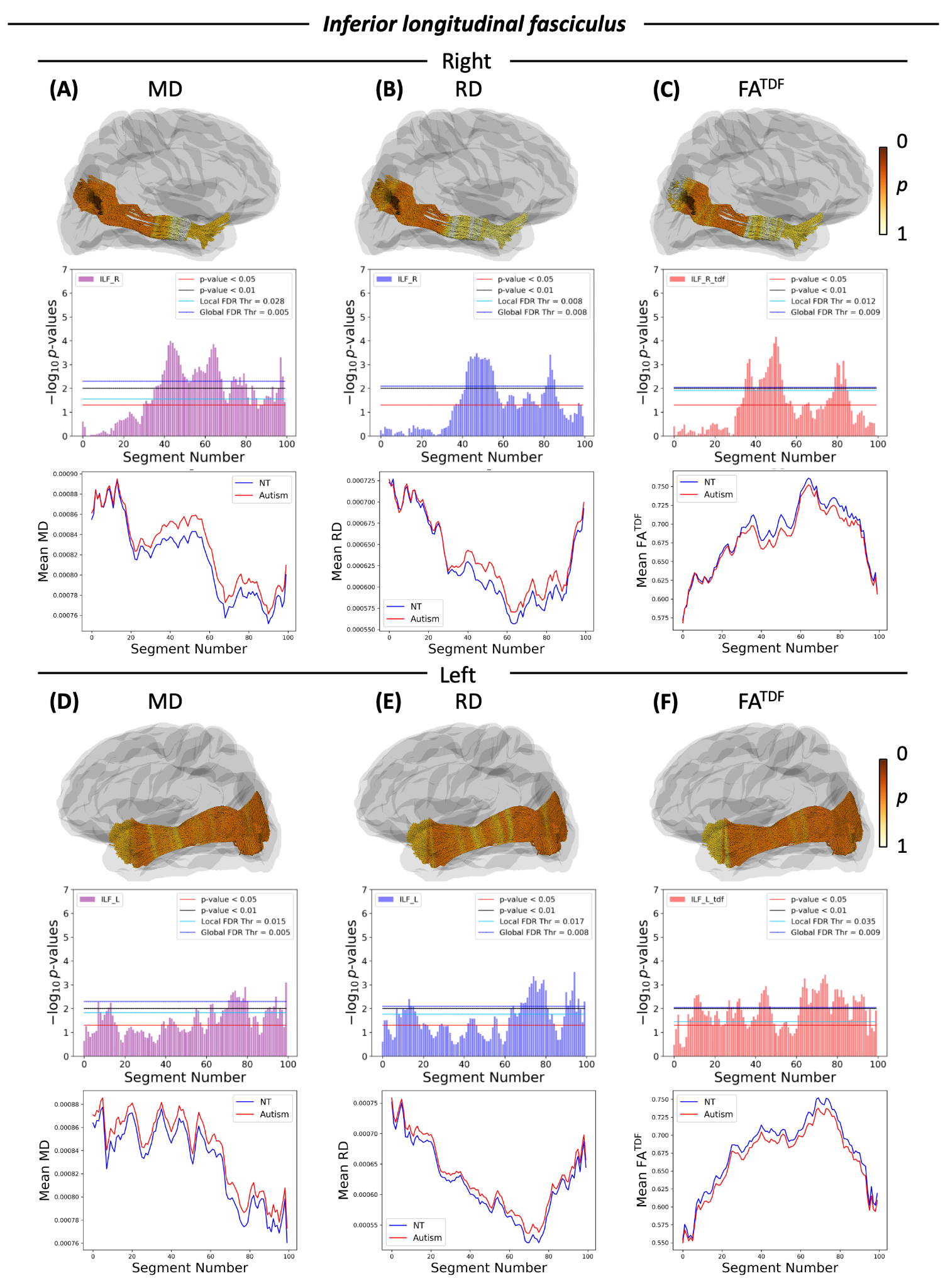

**Supplementary Figure 8.** White matter microstructural alterations in the inferior longitudinal fasciculus in autism compared to neurotypical controls*.* Each column represents a different white matter metric (A) MD, (B) RD, and (C) FA^TDF^ for the right ILF and (D) MD, (E) RD and (F) FA^TDF^ for the left ILF. The first row is a 3D representation of the inferior longitudinal fasciculus illustrating the anatomical location and corresponding *p*-values of the corresponding microstructural metric; the colorbar shows the *p*-values, where darker orange is a lower *p*-value nearing 0 and lighter yellow is a *p*-value of 1. The second row depicts the negative logarithms of *p*-values for each segment along the tract when contrasting the autism and neurotypical groups; the anterior portion of the tract is segment 1, and the segment number increases along the length of the tract up to segment 100 on the posterior portion of the tract. The third row depicts mean plots showing the mean microstructural metric for each group within each segment with autism in red and neurotypicals in blue. FA^DTI^, fractional anisotropy; MD, mean diffusivity; RD, radial diffusivity; FA^TDF^, tensor distribution function fractional anisotropy

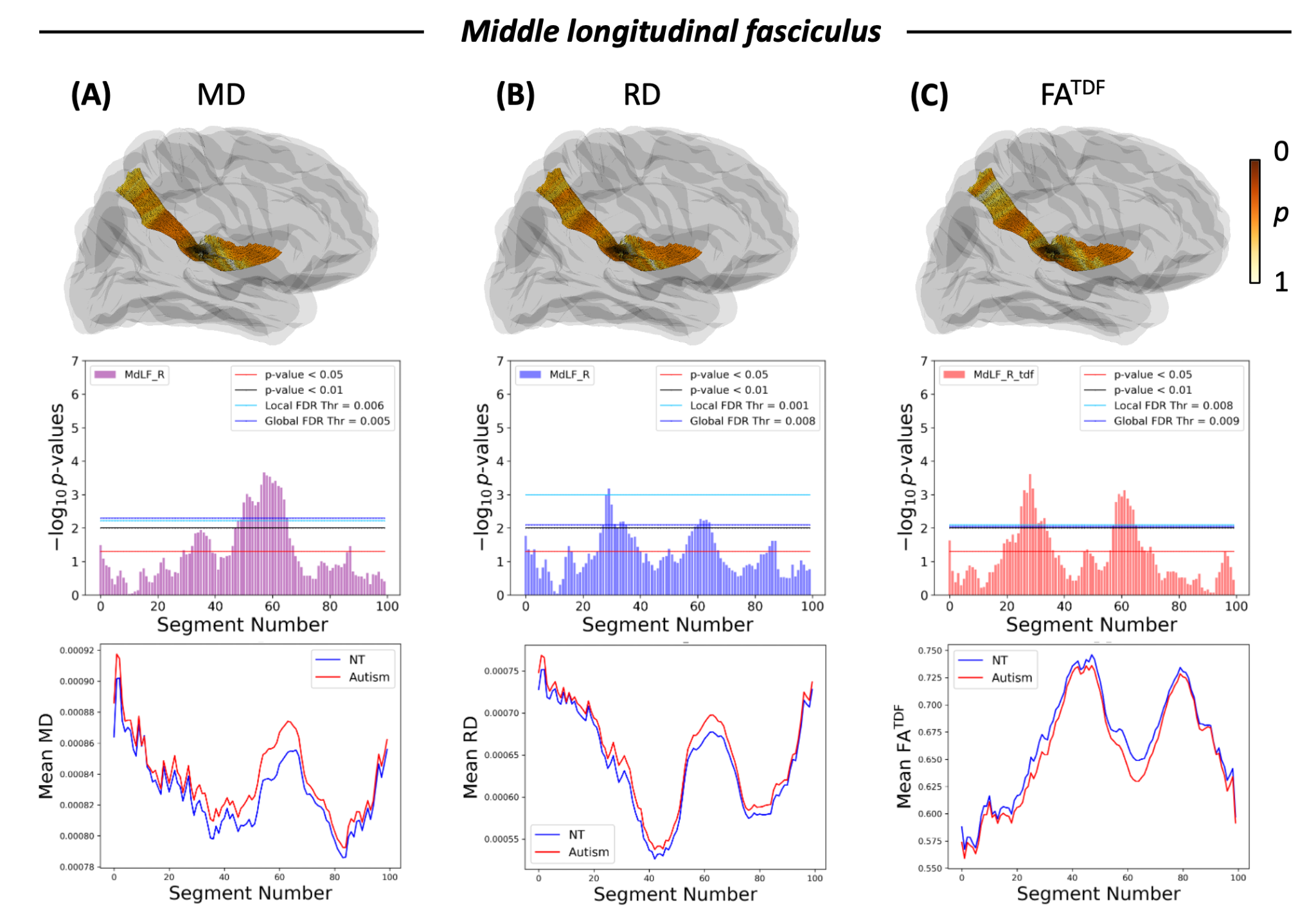

**Supplementary Figure 9.** White matter microstructural alterations in the middle longitudinal fasciculus in autism compared to neurotypical controls*.* Each column represents a different white matter metric (A) MD, (B) RD, and (C) FA^TDF^. The first row is a 3D representation of the middle longitudinal fasciculus illustrating the anatomical location and corresponding *p*-values of the corresponding microstructural metric; the colorbar shows the *p*-values, where darker orange is a lower *p*-value nearing 0 and lighter yellow is a *p*-value of 1. The second row depicts the negative logarithms of *p*-values for each segment along the tract when contrasting the autism and neurotypical groups; the inferior portion of the tract is segment 1, and the segment number increases along the length of the tract up to segment 100 on the superior portion of the tract. The third row depicts mean plots showing the mean microstructural metric for each group within each segment with autism in red and neurotypicals in blue. MD, mean diffusivity; RD, radial diffusivity; FA^TDF^, tensor distribution function fractional anisotropy
